# Attenuating amyloid-beta pathology in mice with in situ programmed astrocytes

**DOI:** 10.1101/2023.10.30.564697

**Authors:** Lun Zhang, Shuai Lu, Ying-bo Jia, Sheng-jie Hou, Jie Zhu, Xiao-ge Liu, Xiao-ying Sun, Ya-ru Huang, Yu-xuan Zhao, Hongan Ren, Chun-yu Liu, Fang Cui, Dong-qun Liu, Xiao-yu Du, Xiao-yun Niu, Ling-jie Li, Ke Wang, Shi-yu Liang, Jin-ju Yang, Shao-yang Ji, Le Sun, Wei-wei Zhou, Xi-xiu Xie, Xiao-lin Yu, Xiaoqun Wang, Rui-tian Liu

**Affiliations:** State Key Laboratory of Biochemical Engineering, Institute of Process Engineering, Chinese Academy of Sciences, Beijing 100190, China; University of Chinese Academy of Science, Beijing 100049, China; State Key Laboratory of Brain and Cognitive Science, CAS Center for Excellence in Brain Science and Intelligence Technology (Shanghai), Institute of Biophysics, Chinese Academy of Sciences (CAS), BNU IDG/McGovern Institute for Brain Research, Beijing 100101, China; Chinese Institute for Brain Research, Beijing 102206, China; Beijing Institute of Brain Disorders, Laboratory of Brain Disorders, Ministry of Science and Technology, Collaborative Innovation Center for Brain Disorders, Capital Medical University, Beijing 100069, China; State Key Laboratory of Primate Biomedical Research, Institute of Primate Translational Medicine, Kunming University of Science and Technology, Kunming, Yunnan 650500, China; Peking-Tsinghua Center for Life Sciences, Academy for Advanced Interdisciplinary Studies, Center for Quantitative Biology (CQB), Peking University, Beijing 100871, China

**Author notes:** These authors equally contributed to this work (Lun Zhang, Shuai Lu, Ying-bo Jia).

## Abstract

Astrocytes are abundant cells in the central nervous system that provide trophic support for neurons and clear detrimental factors, such as Aβ oligomers (AβOs). However, in the brains of Alzheimer’s disease (AD) patients, astrocytes lose these physiological functions. Here, we genetically engineered astrocytes with an anti-AβO chimeric antigen receptor (CAR), constructed by replacing the antigen-binding domain of MerTK with an AβO-specific single-chain variable fragment, to direct their phagocytic activity against AβOs. CAR-engineered astrocytes (CAR-As) showed significantly enhanced phagocytosis of AβOs due to effective activation of Rac1, Cdc42 and RhoA and markedly decreased release of pro-inflammatory cytokines due to inhibition of the NF-κB and cytokine receptor signalling pathways. Consistently, in situ CAR-As markedly ameliorated the cognitive deficits of APP/PS1 transgenic mice possibly by clearing AβOs and creating a non-inflammatory microenvironment for neuronal survival and the restoration of microglia to a healthy phenotype. Our present study is the first to introduce a CAR-A-based therapy, validate its feasibility and effectiveness, and highlight its potential application for the treatment of AD and other brain disorders.

## Introduction

Astrocytes serve a multitude of essential functions in the brain, but the abundant reactive astrocytes in the brains of AD patients lose their normal functions, such as the ability to maintain homeostasis, clear Aβ, and release various detrimental factors, mainly contributing to AD progression(1, 2). Aβ oligomers (AβOs) play a key role in the pathology of AD by inducing neuronal and glial dysfunction and destroying the brain microenvironment(3, 4), contributing a deleterious cycle involving the inability of glia to clear AβOs. Strategies to break this cycle to promote AβO clearance and restore disabled and deleterious astrocytes to a normal phenotype hold great potential to treat AD(5). However, no appropriate astrocyte-targeted therapeutic approach has been reported.

In this study, we proposed a chimeric antigen receptor astrocyte (CAR-A)-based therapeutic strategy for AβO clearance. To avoid the cytokine storm caused by CARs applied in cancer treatment(6), our CAR was constructed based on MerTK, a phagocytic receptor mainly expressed on M2 macrophages and astrocytes(7–9), to allow phagocytosis and simultaneous inhibition of cytokine release upon activation(10, 11). Here, an anti-AβO CAR was constructed by replacing the antigen-binding domain (19Gly-275Asn, Ig-like domain) of MerTK with an AβO-specific single-chain variable fragment, W20(12). When it bound AβOs, the CAR activated the same signalling pathways involved in phagocytosis and anti-inflammation as MerTK, and in situ CAR-engineered astrocytes in mice exhibited significantly enhanced phagocytosis of AβOs and much less inflammatory cytokine release, thus creating a healthy microenvironment in the central nervous system that was beneficial for neuronal survival and glial homeostasis.

## Results

### Anti-AβO CAR expression on astrocytes

To achieve anti-AβO CAR expression on astrocytes, we constructed a plasmid encoding the anti-AβO CAR by replacing the antigen-binding domain of MerTK with W20 and inserting an astrocyte-specific promotor, gfa2(13, 14), adjacent to the W20 gene. The CAR plasmids were loaded in positively charged liposomes conjugated with astrocyte-targeted peptides (AS1, C-LNSSQPS-C)(15) on the surface (Fig. 1a and Fig. S1a, b). Liposome-plasmid complexes (lipoplexes) with an optimal N/P ratio of 4:1 were used for subsequent experiments (Fig. S1c). The size of the lipoplexes was 132.51±19.48 nm, and they had a narrow particle diameter distribution (PDI = 0.19±0.03) (Fig. S1d-f). The MTT results showed that the lipoplexes had minimal toxic effects on astrocytes, neurons and microglia (Fig. S1g-i).

**Fig. 1.**
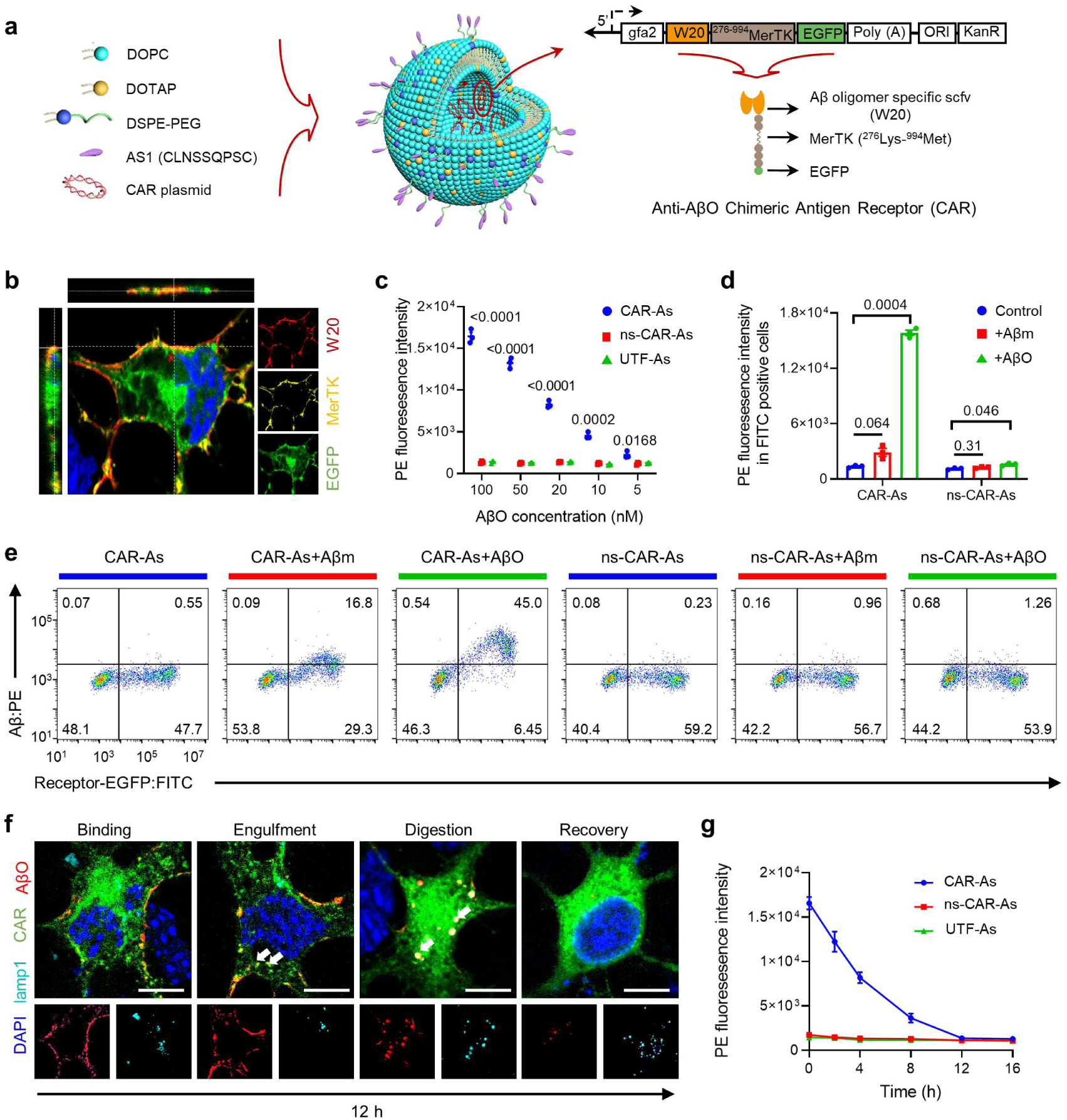
CAR-A generation and their effect on AβO phagocytosis and digestion. **a**, Schematic of astrocyte-targeted lipoplexes used in present study. The designed lipoplexes displays astrocyte-targeted peptides (AS1) on the surface, and contains the plasmids expressing CAR. gfa2, the astrocytes-specific promotor; Poly(A), polyadenylation signal; ORI, origin of replication; KanR, kanamycin resistance gene. **b**, Representative image depicting CAR expression by the co-localization of W20, MerTK and EGFP on astrocytes. Scale bar: 5 μm. **c**, Flow cytometry analysis of the amount of AβOs engulfed by CAR-As, ns-CAR-As and UTF-As in the presence of different AβO concentrations. **d-e**, Flow cytometry analysis of astrocytic engulfment of Aβ monomers and oligomers. The astrocytes were transfected with CAR lipoplexes or ns-CAR lipoplexes for 24 h, after a 6 h-incubation with 200 nM Aβ monomers (Aβm) or Aβ oligomers (AβO), cells were stained with PE-labeled anti-Aβ antibody and analyzed by flow cytometry and PE fluorescence in EGFP positive astrocytes was quantified. **f**, Representative images depicting the phases of engulfment and digestion of AβOs by CAR-As. CAR-As were treated with 200 nM AβOs and washed afterward with fresh medium, Aβ and lamp1 in CAR-As were stained with respective antibody at different times and imaged by confocal microscopy. Scale bars: 5 μm. **g**, The kinetic curves of AβOs digestion in CAR-As, ns-CAR-As and blank astrocytes. For panels **c, d, g,** Data represent the mean ± s.e.m of n = 3 technical replicates and are representative of three experiments. Statistical significance was calculated with one-way ANOVA with multiple comparisons.

To test whether the CARs could be expressed on astrocytes, we transfected primary astrocytes with the prepared lipoplexes and found that CARs expressing W20 (red), ^276-994^MerTK (yellow) and EGFP (green) were successfully expressed on the cell membranes of astrocytes (Fig. 1b) but not neurons or microglia under the control of the gfa2 promotor in the plasmid (Fig. S2a). The transfection efficiency of CAR lipoplexes was 51.4% ± 14.8% in vitro and the expression of CAR on astrocytes lasted for nearly 7 days (Fig. S2b).

### AβO engulfment and digestion by CAR-As

We then evaluated the phagocytic and digestive functions of CAR-engineered astrocytes (CAR-As) against AβOs. The flow cytometry results showed that CAR-As exhibited a significant dose-dependent enhancement of phagocytosis of AβOs but not monomers, while nonspecific scFv-^276-994^MerTK-modified astrocytes (ns-CAR-As) and untransfected astrocytes (UTF-As) showed a marked reduction in the engulfment of Aβ monomers and AβOs (Fig. 1c, d, e) (the small amount of Aβm phagocytosis in Fig. 1d, e was due to the Aβ oligomers aggregated from Aβ monomers in the medium (Fig. S2c)). When blocked with an anti-W20 antibody, CAR-As showed reduced AβO phagocytosis, indicating that W20 mediated the recognition of AβOs by CARs (Fig. S2d). The immunocytochemistry (ICC) results indicated that the process of engulfment and digestion of AβOs by CAR-As could be divided into four consecutive steps: (i) binding, during which AβOs interacted with CARs expressed on astrocytes; (ii) engulfment, during which the AβO and CAR complex was engulfed by astrocytes and phagocytic vesicles were formed; (iii) digestion, during which AβOs were digested through the lysosomal pathway; and (iv) recovery, during which CAR-As were restored to the previous conditions (Fig. 1f, g).

We then explored the signalling pathways involved in AβO engulfment by CAR-As. Only when two adjacent MerTK receptors simultaneously bound to ligands were downstream signalling pathways activated. A single AβO had multiple binding sites for W20, and the binding of two or more CARs on astrocytes to the same AβO was readily induced, resulting in CAR phosphorylation, cytoskeletal remodelling and AβO phagocytosis (Fig. 2a, red box). The phosphorylation of CARs was first confirmed by WB analysis, and the results indicated that CARs were activated and phosphorylated when they interacted with AβOs (Fig. 2b). Consistently, CAR-As showed limited phagocytosis of AβOs when UNC2025, an inhibitor of MerTK phosphorylation, was added (Fig. 2c). It was then revealed by pull-down experiments that downstream Rho family GTPases were activated, and the results showed that Rac1, Cdc42 and RhoA were activated in CAR-As but not in ns-CAR-As or UTF-As during AβO engulfment (Fig. 2d-f), indicating that the assembly of contractile actomyosin filaments and actin polymerization(16) were involved in the engulfment of AβOs by CAR-As.

**Fig. 2.**
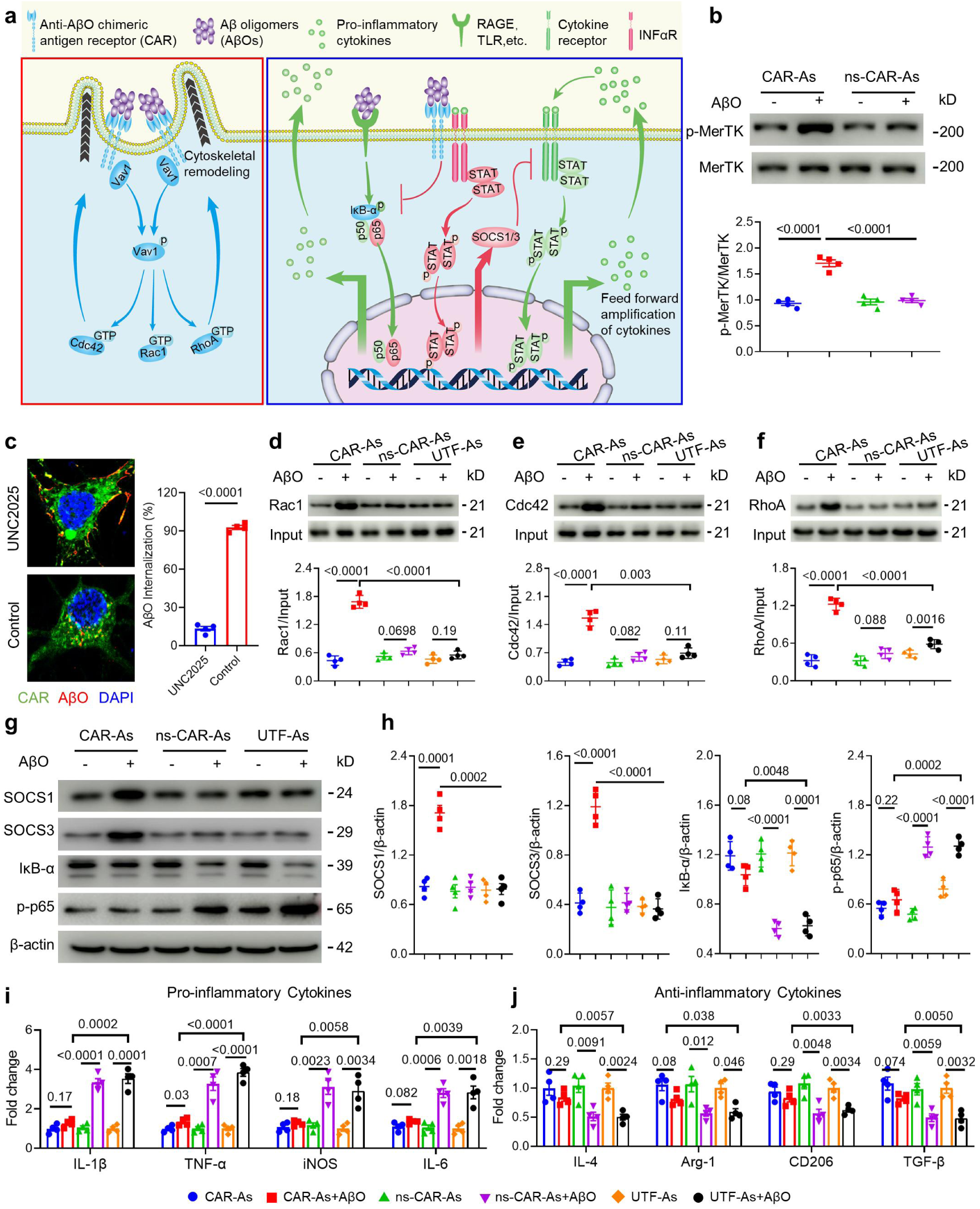
The working mechanism for CAR-As to phagocyte AβOs and modulate microenvironment. **a**, Schematic representation of signaling pathways involved in AβO phagocytosis (red box) and the inhibition of pro-inflammatory cytokine-release (blue box) by CAR-A. **b**, Phosphorylation of CAR or ns-CAR. CAR-As or ns-CAR-As were treated with AβOs for 24 h, the phosphorylation of CAR or ns-CAR was then detected by Western blot, the total CAR or ns-CAR were used as controls. **c**, Left: representative images depicting the effect of UNC2025 on AβOs engulfment by CAR-As. CAR-As were incubated with or without UNC2025 for 12 h, then AβOs were added and incubated for another 6 h. Scale bars: 5 μm. Right: the internalized AβOs in CAR-As was quantified using Image J pro. **d-f**, Activated Rac1 **(d)**, Cdc42 **(e)** and RhoA **(f)** were pulled down from lysates of CAR-As and other control astrocytes treated with or without AβOs for 24 h, and then analyzed by Western blot. Total Rac1, Cdc42 and RhoA were used as controls. Relative levels of activated Rac1, RhoA and Cdc42 were quantified using Image J pro. **g**, The levels of SOCS1, SOCS3, IκB-α and p-p65 in cytoplasm in CAR-As and other control astrocytes treated with or without AβOs for 24 h were analyzed by Western blot, β-actin was used as control. **h**, Relative levels of SOCS1, SOCS3, IκB-α and p-p65 were quantified using Image J pro. The results were combined to express as mean ± s.e.m of four independent repeats. **i-j**, The levels of pro-inflammatory cytokines (IL-1β, TNF-α, iNOS, IL-6) (**i**) and anti-inflammatory cytokines (IL-4, Arg-1, CD206, TGF-β) (**j**) were determined by RT-qPCR. Data are represented as mean ± s.e.m. of n = 3 technical replicates and are representative of four experiments. For all panels, the results were combined to express as mean ± s.e.m of four independent repeats. Statistical significances were determined by one-way ANOVA with Tukey’s multiple comparisons test.

### CAR-As inhibit inflammatory pathways

Mertk activation inhibits the NF-κB pathway(17, 18) and cytokine-receptor cascades(19) (Fig. 2a, blue box). We carried out WB analysis to assay the effect of CAR-As on inflammatory pathways. Consistent with previous reports, the binding of AβOs to advanced glycation end products receptor (RAGE)(20) activated the NF-κB pathway, and inhibitor of κB α (IκB-α) degradation and increased phosphorylation of p65 (p-p65) levels were observed of UTF-As and ns-CAR-As. However, the binding of AβOs to CAR-As inhibited the NF-κB pathway, and IκB-α levels and p-p65 levels were not significantly changed (Fig. 2g, h). Moreover, the binding of AβOs to CAR-As enhanced the expression of suppressor of cytokine signalling protein 1/3 (SOCS1/3)(21) (Fig. 2g, h), thus inhibiting the feed-forward amplification of cytokines by reducing the release of pro-inflammatory cytokines, including IL-1β, TNF-α, iNOS, and IL-6 (Fig. 2i), and increasing the generation of anti-inflammatory cytokines, including IL-4, Arg-1, CD206, and TGF-β (Fig. 2j). The levels of pro- and anti-inflammatory cytokines were confirmed by enzyme-linked immunosorbent assay (ELISA) (Fig. S2e).

### CAR-As improve the microenvironment

Microenvironmental changes in the brain are vital for the pathogenesis of AD(22). To test the influences of CAR-As on the microenvironment and other cells during AβO clearance, we added conditioned medium (CM) from CAR-As and other experimental cells to cultured microglia and neurons (Fig. S3a). CM from CAR-As but not from ns-CAR-As or UTF-As significantly decreased pro-inflammatory cytokine generation, restored anti-inflammatory cytokine expression in cultured microglia (Fig. S3b-f), and decreased the detrimental effects of AβOs on neurons by maintaining the length of axons and dendrites (Fig. S3h, i). The neutralizing antibodies to pro-inflammatory cytokines or anti-inflammatory cytokines were then used to identify the dominant cytokines that influenced the microenvironment. The results showed that multiple cytokines rather than single cytokine affected the microenvironment (Fig. S3j).

### Efficient generation of CAR-As in vivo

In vivo, we first evaluated the effectiveness of CAR lipoplexes delivery to the brain and the ability of CAR lipoplexes to infect cells when delivered by intranasal (I.N.) administration, a non-invasive delivery option. The distribution of the CAR lipoplexes labelled with CY-7 in mice was visualized after I.N. administration, and the results demonstrated that the CAR lipoplexes were mainly localized in the brain, lungs and livers and most of them were metabolized within 24 hours (Fig. S4a, b). Haematoxylin and eosin (H&E) staining and histology score(23) of the brains, hearts, livers, spleens, lungs, kidneys, pancreases, skin and bone revealed no cellular morphological change in the organs of mice treated with CAR lipoplexes or ns-CAR lipoplexes (Fig. S4c, d).

The IHC results showed that CARs were mainly expressed in astrocytes in the brain (Fig. S5a), and the flow cytometry analysis showed that the transfection efficiency of astrocytes in the hippocampus and cortex was 5.03±2.17% and 3.70±1.78%, respectively (Fig. S5b, h). In order to determine the transfection specificity, we sorted out the cells that were positively transfected in the brain by FACS and measured the expression of astrocyte marker genes (*Gfap, Aqp4*) by qPCR, and the results further confirmed that the positively transfected cells were astrocytes (Fig. S5c-e). Furthermore, IHC and flow cytometry analysis showed that, except for brain, there was no expression of CARs in other tissues and organs (Fig. S5f-g), which should be attributed to the gfa2 promotor and AS1 peptide in the CAR lipoplexes.

### CAR-As attenuate cognitive deficits

We next tested the therapeutic effects of CAR-As on a transgenic AD-like mouse model (APP/PS1). The mice were intranasally treated with saline, empty liposomes, CAR lipoplexes, or ns-CAR lipoplexes for six weeks (Fig. S6a). No significant differences in mouse survival, body weight, or motor function were observed (Fig. S6b-d). A water maze experiment was then conducted to assess the spatial learning and memory of the mice. During the training phase, mice treated with CAR lipoplexes displayed significantly improved spatial learning, showing shorter latency time than saline-, empty liposomes-, and ns-CAR lipoplexes-treated APP/PS1 mice (Fig. 3a,b). A probe test in which the platform was removed was then conducted to evaluate memory recall. Consistently, retention memory was significantly improved in CAR lipoplexes-treated mice, as they exhibited shorter latency time, spent a longer time in the target quadrant and crossed the platform more times than the mice in the other groups (Fig. 3c-e).

**Fig. 3.**
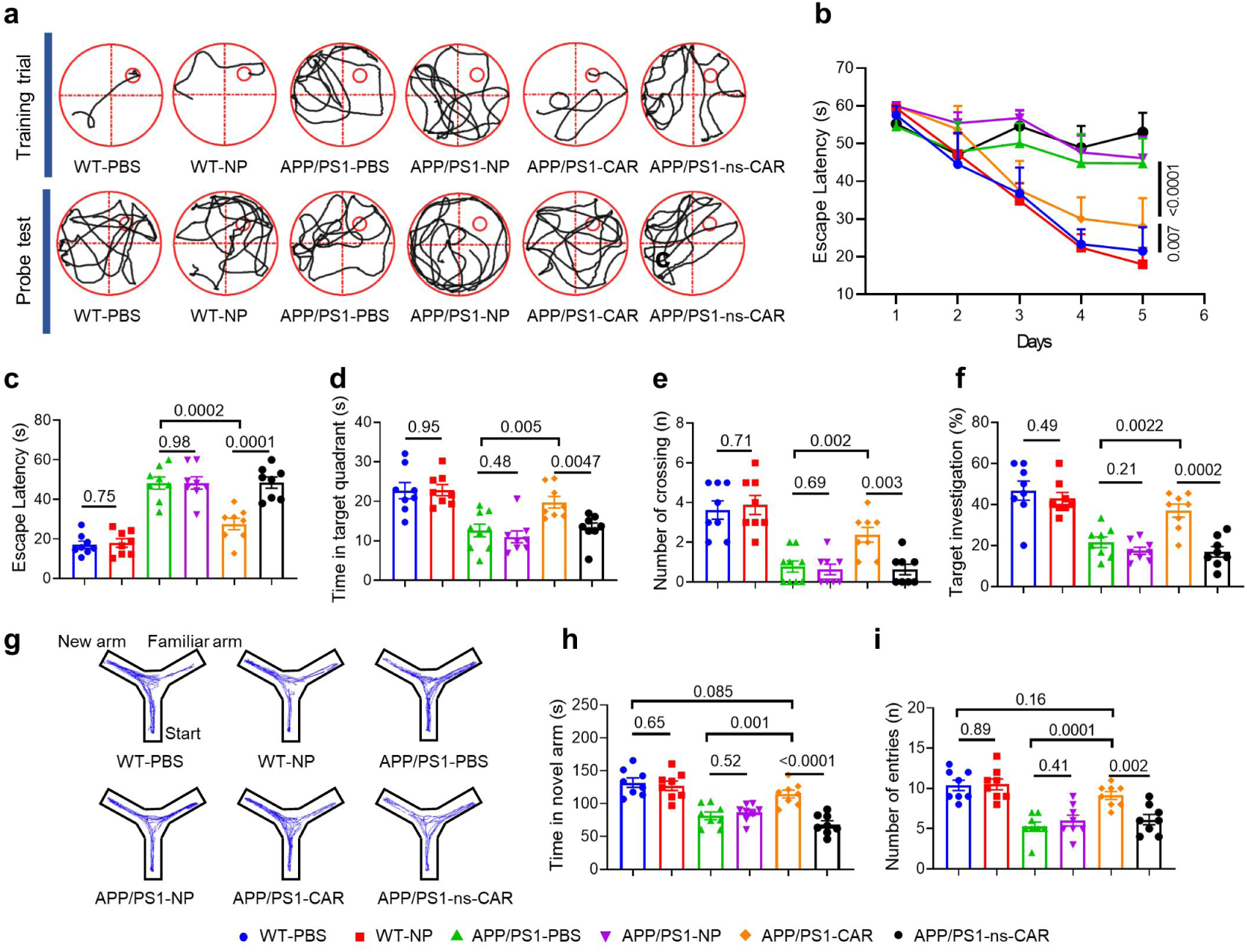
CAR-As attenuated memory deficits in APP/PS1 mice. Wild type mice were intranasally treated with saline and empty liposomes, and APP/PS1 mice were treated with saline, empty liposomes, CAR lipoplexes and ns-CAR lipoplexes for 6 weeks. **a**, Representative swimming paths in training trial (up) and probe test (down) in Morris water maze. **b**, The time needed to reach the hidden platform was plotted across training days (*p*-value represented the comparison of APP/PS1-CAR group with APP/PS1-PBS and WT-PBS, respectively). **c-e**, The latency to find the position of the platform (**c**), the time spent by the mice in the targeted quadrant (**d**) and the number of platform crossings (**e**) were recorded during the probe test after the training phase with the platform removed. **f**, The percentage of target investigation in novel object recognition test in different experimental groups. **g**, Representative tracks in Y maze test for different experimental groups. **h-i**, The time spent in the novel arm (**h**) and the number of entries (**i**) in Y-maze test for different experimental groups. For panels **c-f** and **h-i**, the experiment was performed once. Values are the mean ± s.e.m. of n = 6-8 mice per group. A one-way ANOVA with post hoc Tukey test was used for statistical analysis.

To further evaluate the hippocampus-dependent spatial working memory and short- and long-term recognition memory of mice, the Y-maze and novel object recognition (NOR) tests were conducted. The results showed that CAR lipoplexes-treated mice exhibited a significant increase in exploration of the novel object in the NOR test (Fig. 3f) and spent more time in and made more entries into the novel arm of the Y-maze (Fig. 3g-i). These results indicated that the cognitive and memory deficits was relieved in CAR-A treated APP/PS1 mice.

### CAR-As decrease Aβ load

To assess the effect of CAR-As on the level of AβOs in APP/PS1 mouse brains, immunohistochemistry (IHC) was conducted. The results demonstrated that unlike the other treatments, CAR lipoplexes treatment greatly enhanced AβO engulfment by CAR-As (Fig. 4a-c), and the proportion of the CAR-As that engulfed AβO to CAR-As was 54.2 ± 17.8% and 58.6 ± 12.3% in hippocampus and cortex, respectively. Consistently, the levels of AβOs in brain homogenates (Fig. 4d) and brain interstitial fluid (ISF) (Fig. 4e), Aβ42, Aβ40 and plaques were reduced in the brains of mice treated with CAR lipoplexes but not those treated with saline, empty liposomes, or ns-CAR lipoplexes (Fig. 4f-i). Moreover, compared with the APP/PS1-PBS and APP/PS1-ns-CAR groups, the Aβ levels in the cerebrospinal fluid but not in the serum of the APP/PS1-CAR group decreased to a certain extent (Fig. S7a).

**Fig. 4.**
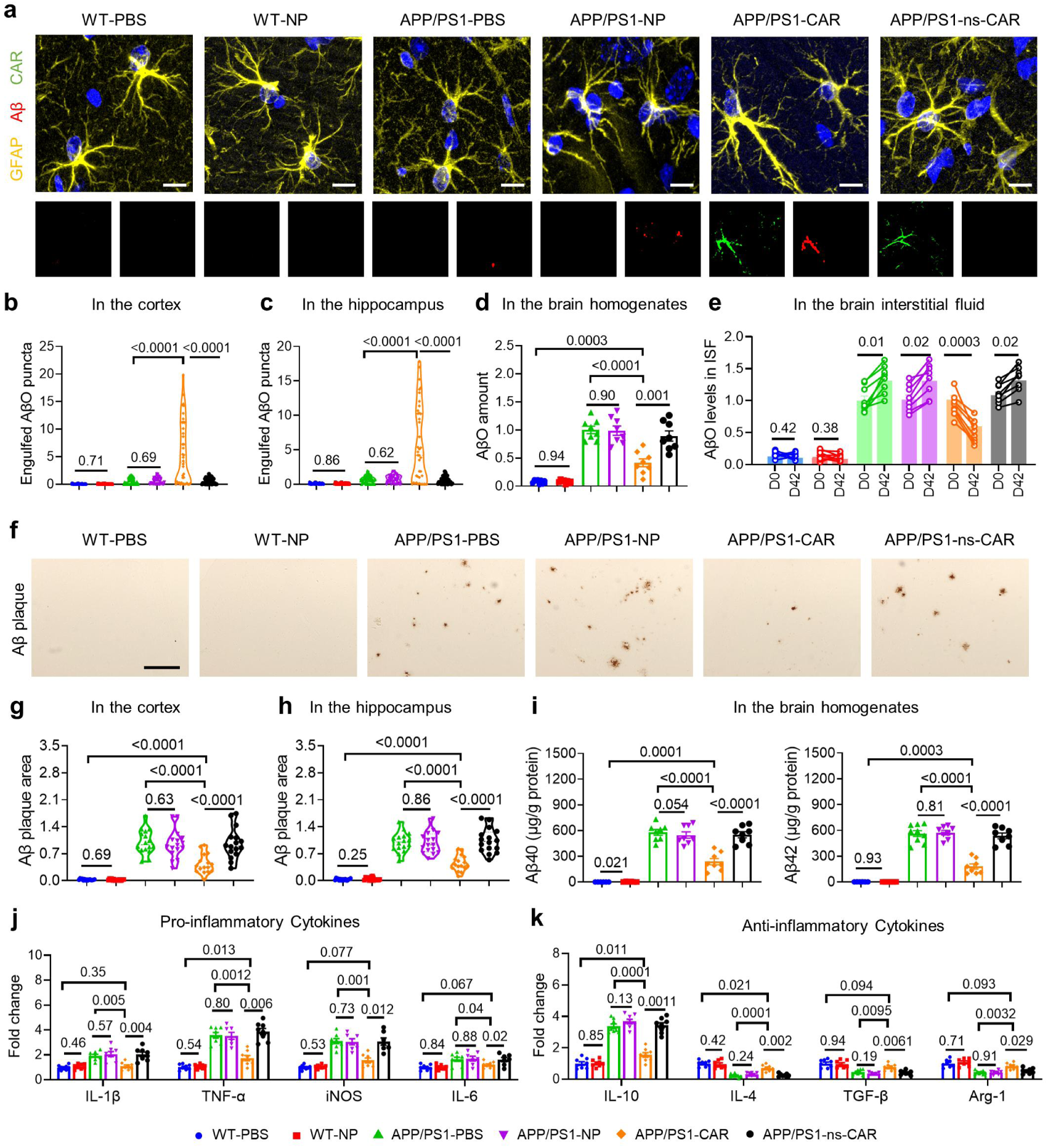
CAR-As attenuated the neuropathology in APP/PS1 mouse model. Wild type mice were intranasally treated with saline and empty liposomes, and APP/PS1 mice were treated with saline, empty liposomes, CAR lipoplexes and ns-CAR lipoplexes for 6 weeks. **a**, Representative images depicting AβOs engulfment by different experimental groups. Scale bars: 5 μm. **b-c**, the amount of AβOs in astrocytes in cortex (**b**) and hippocampus (**c**) were quantified using Image J pro. Values were normalized against the APP/PS1-PBS group. **d**, The amount of AβOs in mouse brains in different experimental groups was quantified by ELISA. Values were normalized against the APP/PS1-PBS group. **e**. AβO levels in brain interstitial fluid (ISF) of the same mouse in different experimental groups at day0 (D0) and day42 (D42) was determined by microdialysis. Values were normalized against the APP/PS1-PBS group at D0. **f**, Representative images of Aβ plaques in mouse brains in different experimental groups. Scale bar: 200 μm. **g-h**, the area of Aβ plaque in cortex and hippocampus were quantified using Image J pro. Values were normalized against the APP/PS1-PBS group. **i**, Aβ40 and Aβ42 in mouse brains in different experimental groups were measured by ELISA. **j-k**, The levels of the pro-inflammatory cytokines (IL-1β, TNF-α, iNOS, IL-6) (**j**) and anti-inflammatory cytokines (IL-10, IL-4, Arg-1, TGF-β) (**k**) in mouse brains from different experimental groups were determined by RT-qPCR. For panels **b-c** and **g-h**, Data are represented as the mean ± s.e.m. of n = 40. A one-way ANOVA with post hoc Tukey test was used for statistical analysis. For panels **d-e** and **i-k**, Data are represented as the mean ± s.e.m. of n = 6-8 mice per group. A one-way ANOVA with post hoc Tukey test was used for statistical analysis.

### CAR-As improve brain microenvironment

The expression of pro-inflammatory and anti-inflammatory cytokines in the mouse brain was measured by quantitative reverse transcription PCR (RT-qPCR). The results revealed that the expression of pro-inflammatory cytokines was significantly reduced and that the expression of anti-inflammatory cytokines was upregulated in the brains of CAR lipoplexes-treated mice compared with those of saline-, empty liposomes-, and ns-CAR lipoplexes-treated APP/PS1 mice (Fig. 4j, k), which was consistent with the protein levels determined by ELISA (Fig. S7b, c). Moreover, our IHC and WB results showed significantly reduced microgliosis and astrogliosis in the brains of CAR lipoplexes-treated mice compared with the brains of mice in other groups (Fig. S7d-f). These data indicated that CAR-As relieved neuroinflammation and improved the brain microenvironment in APP/PS1 mice.

### The expression profiles of CAR-As

RNA-seq analysis was performed to analysis gene transcriptional profiles of astrocytes isolated from wildtype mouse (astrocyte^WT^) and APP/PS1 mouse (astrocyte^APP/PS1^), CAR positive astrocytes (CAR-A^+^) and CAR negative astrocytes (CAR-A^-^) isolated from APP/PS1-CAR group. Compared with the astrocyte^APP/PS1^, CAR-A^+^ cells in APP/PS1 mice significantly reduced expression of inflammation-related genes and increased the expression of phagocytic function-related genes (Fig. S8a). Interestingly, CAR-A^-^ cells changed the expression of these genes to some extent still in a similar way as CAR-A^+^ cells, indicating that CAR-A^+^ affected neighboring astrocytes (Fig. S8b).

### CAR-As regulate microglial biofunction

Microglia are highly plastic cells that change their phenotype and biological characteristics in response to the microenvironment(24). To test the effect of CAR-As on the phenotype of microglia in the APP/PS1 mouse brain, microglia were isolated from the brains of mice subjected to various treatment, and the expression levels of associated genes were measured. Microglia from CAR lipoplexes-treated APP/PS1 mice switched to a healthy phenotype, as indicated by expression of M0 microglia-associated genes, while microglia from saline-, empty liposomes-, ns-CAR lipoplexes-treated APP/PS1 mice maintained the MGnD phenotype (Fig. S9a, b). WB analysis of the protein levels of P2ry12 and Clec7a revealed consistent results (Fig. S9c). To evaluate the lysosomal function of microglia in the APP/PS1 mouse brain, the expression of CST7(25), a lysosomal inhibition protein, was measured. WB analysis demonstrated that the levels of CST7 were significantly lower in microglia from CAR lipoplexes-treated APP/PS1 mice than in those from control APP/PS1 mice (Fig. S9c), indicating that lysosomal function of microglia was partially restored in CAR lipoplexes-treated APP/PS1 mice. Moreover, TNF-α, TGF-β(*1*) and two other key cytokines TGF-α, VEGF-β(26) secreted by microglia that regulates astrocyte activation were measured, and the results indicated that these cytokines were apt to return to normal levels in microglia from CAR lipoplexes-treated APP/PS1 mice (Fig. S9d).

### CAR-As improve the number of synapses

Synapse damage is another pathological feature of AD(27). Here, we determined the levels of postsynaptic density-95 (PSD95) and synaptophysin (SYN) in the brains of APP/PS1 mice from the different groups. The results showed that the levels of PSD95 and SYN and the number of intact synapses (indicated by co-localization of PSD95 and SYN) were significantly enhanced in the CAR lipoplexes-treated group compared with the saline-, empty liposomes-, and ns-CAR lipoplexes-treated APP/PS1 mice (Fig. S9f, g), and these findings was further confirmed by WB analysis of synapse marker protein levels (Fig. S9h). Moreover, we measured the changes of two neurotransmitters, glutamate and acetylcholine in the brains of mice in different experimental groups, and found that CAR lipoplexes treatment partially restored the levels of the two neurotransmitters (Fig. S9e).

### CAR-As retain major astrocytic biofunctions

We next characterized astrocytic changes pre- and post-transfection in vitro and in vivo through series of experiments. The morphology of CAR-A and normal astrocytes in vivo were measured by confocal imaging, and no significant change between CAR-As and normal astrocytes was observed (Fig. S10a, b). The important proteins in the CNS secreted from astrocytes such as amyloid precursor protein (APP), apolipoprotein E (APOE) and thrombospondin 1 (TSP1) did not significantly changed between CAR-As and UTF-As (Fig. S10c, d). The glutamate uptake ability of CAR-As was also measured, and no significant change between CAR-As and normal astrocytes was observed (Fig. S10e). The conditioned medium from CAR-A and UTF-A showed no difference in neuron trophic support (Fig. S10f), synaptogenesis and synapse maturation (Fig. S10g), and synapse function (Fig. S10h-j). CAR-A and UTF-A also showed no difference in engulfment of myelin debris and synaptosomes (Fig. S10k). To detect the effect of CAR-A on BBB integrity, we examined the infiltration of Evans blue (EB) dye in the brains of wildtype mice and CAR-A mice, and we did not find the significant influence of CAR-A on BBB integrity (Fig. S10l). A normal karyotype of CAR-A including 19 pairs of autosomes and a pair of sex chromosomes (XY) was observed (Fig. S11a). Moreover, RNA-sequencing (RNA-seq) analysis indicated that the genes implicated in receptor internalization (Mx1, Mx2), protein phosphorylation (Mylk3, Lama1), leucyl-tRNA aminoacylation (Lars2), and CAR expression (Mertk) were upregulated, and the genes implicated in negative regulation of transcription (Scml2), G protein-coupled receptor signaling pathway (Olfr539) were downregulated (Fig. S11b). CAR transfection did not change significantly the expression in astrocyte makers, cholesterol metabolism, synapse maintenance and immune/antigenic response (Fig. S11c). Furthermore, a gene set enrichment analysis on 56 strongest differentially expressed genes between WT astrocyte and CAR-A was performed and the top significant gene sets are showed in Fig. S11d. The gene set related to immune response and nucleotide binding events showed significant enrichment in the astrocyte versus CAR-A group.

## Discussion

AβOs play an important role in AD progression by receptor binding, cell membrane destruction, mitochondrial damage, Ca^2+^ homeostasis dysregulation, tau pathological induction and neuronal death. Moreover, AβOs can induce microglia to transform into MGnD phenotype, which releases inflammatory factors and further induces neuronal death. Therefore, effectively removing AβOs can reduce the levels of AβOs and inflammatory factors, and prevent neuronal death, normalizing brain microenvironment, reversing MGnD microglial phenotype and restoring the number of synapses(28). CAR-A technology is an ideal strategy to endow astrocytes with the ability to engulf AβOs and simultaneously block neurotoxin generation, which is achieved by the binding of two CARs to AβOs and inhibition of the NF-κB pathway, creating a healthy microenvironment beneficial for neurons and other neural cells.

To improve the translational application of our strategy, we chose liposome nanoparticles, which have been extensively used clinically with less safety concerns, to deliver the CAR plasmid, and obtained an ideal astrocyte transformation efficiency and consistent expression of CAR on astrocytes by convenient nasal spray twice per week. Compared with microglia, astrocytes are easy to transfect besides of their number advantage. The CAR can be expressed and displayed on astrocytes for 5-7 days to mediate the phagocytosis of Aβ oligomers in mouse brain, which was consistent with the CAR expression in vitro.

CAR-based strategy was an effective therapy for APP/PS1 mice that did not cause observable side effects, and CAR-As themselves retained multiple astrocytic biofunctions. Additionally, the increased acetylcholine levels in the mouse brains were induced by the recovery of the physiological function of astrocytes post CAR treatment, which were significantly higher than that in APP/PS1 mice, but they did not exceed that in WT mice and were within physiological range. Therefore, CAR-A therapy is unlikely to cause Parkinson’s disease-like neuropathology. These results suggest that CAR-As expressing different scFv antibodies hold beneficial potential to treat a variety of chronic neurological diseases without observable side effects. Although the engulfment of pathogenic agents is not the main function of astrocytes, the CAR-A-based technology improved the phagocytic ability of astrocytes and exerted therapeutic effects; thus, it provides new ideas for the functional modification of other types of glial cells for the treatment of AD and other brain disorders.

## Methods

### Primary microglial and astroglial cultures

Primary astrocytes and microglia cultures were prepared from APP/PS1 mice as described previously(29). Briefly, the brain cortices from 10 postnatal mice were dissected in Dulbecco’s PBS (dPBS) and the meninges were removed. The tissues were enzymatically dissociated with papain enzyme at 5% CO_2_ and then mechanically dissociated to produce single cell suspension. For astrocyte isolation, 100 μl of 0.5 mg/ml sheep anti-ITGB5 (R&D Systems, AF3824) was added into 5-10 ml of cell suspension and incubated with the cells for 30-40 min at 37°C, and cell suspensions were allowed to interact with the immunopanning dish (coated with 60 μl of secondary antibodies in 50 mM Tris-HCl pH 9.5 overnight at 4°C) for 20 min at room temperature, unbound cells and debris were removed by washing the dish with dPBS ten consecutive times, freshly isolated astrocytes were cultured in DMEM/Neurobasal (1:1) containing 100 units/mL penicillin, 100 μg/mL streptomycin (ThermoFisher Scientific,15140122), 1 mM sodium pyruvate (ThermoFisher Scientific, 11360070), 292 μg/ml L-glutamine (ThermoFisher Scientific, 25030081), 1 μg/mL transferrin (Sigma, T8158), 0.16 μg/mL putrescine (Sigma, P5780), 1 nM progesterone (Merck, P0130), 0.4 ng/ml sodium selenite (ThermoFisher Scientific, 11360070), 5 μg/ml N-Acetyl-L-cysteine (Sigma, A8199) and 5ng/ml HBEGF (MedChemExpress, #HY-P7400). For microglia isolation, cell suspensions were applied directly to positive-selection immunopanning dishes coated with anti-CD11b monoclonal antibodies, freshly isolated microglia were cultured in DMEM/F12 containing 100 units/mL penicillin, 100 μg/mL streptomycin, 2 mM L-glutamine, 5 μg/ml N-acetyl cysteine, 5 μg/ml insulin (Merck, I9278), 100 μg/mL transferrin, and 100 ng/mL sodium selenite.

### Primary neuronal cultures

Primary neurons were obtained from newborn pups of APP/PS1 mice. Briefly, hippocampi were dissected out from the brains and incubated with trypsin and DNase I before careful trituration to generate single-cell suspensions. The cells were plated on poly-D-lysine coated coverslips at a density of 300,000/well in 12-well dish and cultured in neurobasal medium with B27and L-GlutaMAX. The medium was replaced every 2-3 days.

### Plasmid construction

All of the plasmids used in this project were custom-cloned by Sangon Biotech (Shanghai) Co., Ltd. The following 2 gene expression vectors were used:

(1) pgfa2-W20-^276-994^MerTK-EGFP-BGH polyA

In this construct, enhanced CMV promotor sequence in MerTK expression plasmid (purchased from Sino Biological Inc., #MG50514-ACG) was replaced with gfa2 promotor sequence^14^ synthesized from Sangon. The antigen-binding domain gene corresponding to amino acids 19-275 of MerTK was replaced with W20 (AβOs-specific single-chain variable fragment) gene sequence(12). To facilitate the detection of receptor, we co-expressed enhanced green fluorescent protein (EGFP) along with the ^276-994^MerTK before the stop codon and the bovine growth hormone (BGH) poly-A signal.

(2) pgfa2-ns-scFv-^276-994^MerTK-EGFP-BGH polyA

In this construct, W20 single-chain variable fragment gene sequence in plasmid (1) was replaced with the an nonspecific single-chain variable fragment (ns-scFv) gene sequence(30).

### Synthesis of DSPE-PEG-AS1

AS1 peptides (synthesized from Chinese Peptide Company) were conjugated with DSPE-PEG2000-NHS (Aladdin, #D163641) (1.5:1 molar ratio) in N,N-dimethylformamide containing triethylamine (5 μL mL^-1^) at room temperature for 72 h under gentle stirring, the resulting reaction solution was dialyzed against distilled water (cutoff of molecular weight=2000 Da) for 48 h. The final solution after dialysis was freeze-dried and stored at -20°C. The conjugations were confirmed by determining the molecular weight of the resulting DSPE-PEG2000-AS1 using a MALDI-TOF mass spectrometer.

### Preparation of Aβ42 oligomers

Aβ42 peptide (synthesized from Chinese Peptide Company) was dissolved in PBS at 100 μM and incubated at 37 °C without agitation for 4-8 h, and then separated by size exclusion chromatography.

### Preparation and characterization of liposomes and lipoplexes

DOTAP (Avanti Polar Lipids, #890890), DOPC (Aladdin, #D130438), and DSPE-PEG2000-AS1 were dissolved in chloroform/methanol mixed solution in a 2:1 ratio at a concentration of 9.5, 9.5, and 1 μmol mL^-1^, respectively. After gentle stirring at room temperature for 2 h, the mixture was rotated on a rotary evaporator at 40 °C to make a thin film, the dried lipid film was then hydrated using 10 mM Tris-buffer (pH 7.4) followed by sonicating using a probe sonicator for two 5 min periods with an interval of 1 min in an ice-water bath to generate small unilamellar vesicles (SUVs) liposomes. The resulting liposomes were extruded five times through a 200 nm polycarbonate membrane and five times through a 100 nm membrane (Avanti Polar Lipids) using an Avanti Mini Extruder (Avanti Polar Lipids). For preparation of plasmid-liposome complexes (lipoplexes), mixture of plasmids and liposomes at different ratio was vortexed and kept at room temperature for 15-20 min. Then, the prepared lipoplexes were transferred into a cryogenic vial to perform ten freeze-thaw cycles, followed by extruded five times through 200 nm and 100 nm membranes.

Optimal N:P ratios of lipoplexes was determined by gel retardation assay using 1% agarose gel (Solarbio, #A8201) containing 0.5 μg mL^-1^ ethidium bromide (Thermo Fisher Scientific, #17898). TEM was used for the morphological examination of the nanoparticles operating at an accelerating voltage of 100 kV (Hitachi, HT7700). The hydrodynamic sizes and zeta potentials of the liposomes and lipoplexes were characterized at room temperature and at a scattering angle of 90° by differential light scattering using Zetasizer Nano ZSP (Malvern).

MTT assay was performed to assess the cytotoxicity of liposomes and lipoplexes on astroglia, microglia and neurons. In brief, the cells were seeded in 96-well plates with approximately 10^4^ cells per 100 μL of medium per well. The plates were incubated at 37 °C for 24 h to allow the cells to attach. The samples at different concentrations were added to individual wells. The plates were incubated for another 48 h at 37 °C. The medium was then replaced with 100 μL DMSO to dissolve the crystals formed in the previous step and then incubated for 2 hours at room temperature in the dark. Finally, absorbance was measured spectrophotometrically at 570 nm using a SpectraMax M5 spectrophotometer (Molecular Devices).

### Flow cytometry and cell sorting

CAR-As with CARs expression and AβOs phagocytosis were tested by flow cytometry using a two-step staining protocol(31). Briefly, after fixed in 4% PFA for 20 min, transfected astroglia were stained with anti-Aβ primary antibody, followed by anti-mouse PE (Biolegend, #406607) secondary staining. Flow cytometry data were acquired on a CytoFLEX LX (Beckman) and analyzed with FlowJo X10 (FlowJo). For experiments involving CAR-As sorting, CAR-positive astroglia were sorted using a FACSAria II cell sorter (BD Biosciences).

### Adult astrocyte isolation

Mice were perfused with 0.9% saline under isoflurane anesthesia, and then the brain was removed, dissected, and rinsed in HBSS. Enzymatic cell dissociation was then performed using an Adult Brain Dissociation Kit (130-107-677, Miltenyi Biotec), according to the manufacturer’s instructions. The resulting single cell suspension was centrifuged at 300g for 10 min at RT, resuspended in 40% Percoll and centrifuged at 800×g for 20 min with breaks off. After Fc receptors were blocked using anti-mouse CD16/CD32 (1:100, eBioscience, 14-0160-82) for 10 min, cells were incubated with ACSA-2-PE for 30 min on ice, washed with 1 ml of blocking buffer and sorted using a FACSAria II cell sorter.

### RNA-seq and data analysis

Total RNA was isolated from astrocytes of mouse brain using a RNeasy mini kit (Qiagen). For each sample, 1 μg total RNA was used for library preparation. In brief, the poly(A) mRNA isolation was performed using Oligo(dT) beads and the mRNA fragmentation was performed using divalent cations and high temperature. Priming was performed using Random Primers. First strand cDNA and the second-strand cDNA were synthesized. The purified double-stranded cDNA was then treated to repair both ends and add a dA-tailing in one reaction, followed by a T-A ligation to add adaptors to both ends. Size selection of adaptor-ligated DNA was then performed using DNA Clean Beads. Each sample was then amplified by PCR using P5 and P7 primers and the PCR products were validated. Then libraries with different indexs were multiplexed and loaded on an Illumina Novaseq instrument for sequencing using a 2×150 paired-end configuration according to manufacturer’s instructions. DEseq was used to identify differentially expressed genes (DEGs) and the significant genes with fold change>2, and adjusted p value<0.05 were used for subsequent analysis and hierarchical clustering was performed as described(32).

### Adult microglia isolation

Primary microglia were isolated from the mice brain as previously described^40^. In brief, the brains of mice were minced in the Hibernate A (Thermo Fisher Scientific, #A1247501)/B27 medium and dissociated for 15 minutes at 37 °C with 0.25% Trypsin-EDTA containing 1 mg mL^-1^ DNase I, after neutralized with DMEM supplemented with 10% FBS, cells were separated by Optiprep (Merck, #D1556) density gradient centrifugation. Fractionated microglia from APP/PS1 and WT mice were obtained for further quantitative PCR and Western blot assays.

### Quantitative RT-qPCR analysis

RNA was extracted from cell lysates and brain homogenates using the RNeasy Lipid Tissue kit (Qiagen, #74804) according to the manufacturer’s instructions. cDNA was prepared from total RNA using the PrimeScript RT-PCR kit (Takara, #RR037Q). Relative gene expression of the cDNA was assayed using a 7500 Fast real-time PCR instrument (Applied Biosystems) with SYBR Select Master Mix (Applied Biosystems, #4472908), qPCR data were analyzed by the ΔΔC_T_ method by normalizing the expression of each gene to housekeeping gene GAPDH and then to the control groups. The used primers in this study are listed in Supplementary Table 1.

### Pull-down Assay

Active RhoA and active Cdc42/Rac1 were pulled-down by GST-Rok (RBD) and GST-Pak (PBD) beads according to the previous method. Briefly, RBD and PBD fusion proteins were expressed in BL21(DE3) bacteria and captured on Glutathione Sepharose 4B (Cytiva, # 17075601), protein lysates of CAR-As, ns-CAR-A and blank astrocytes treated with or without 200 nM AβOs for 24 h were mixed with 20 μL of PBD or RBD beads and incubated at 4 °C for 45 min. Then, the beads were washed and the proteins bound on the beads were analyzed by Western blotting.

### Electrophysiology

Whole-cell recording was performed on cultured primary neurons at room temperature. Randomly selected neurons held at -70mV and mEPSC were recorded in oxygenated (95% O_2_ and 5% CO_2_) recording solution (in mM: 126 NaCl, 3 KCl, 26 NaHCO_3_, 1.2 NaH_2_PO_4_, 10 D-glucose, 2.4 CaCl_2_, 1.3 MgCl_2_ and 1 µM TTX). Patch pipettes had a 6-10 MΩ resistance when filled with intracellular solution (in mM: 130 potassium gluconate, 16 KCl, 2 MgCl_2_, 10 HEPES, 0.2 EGTA, 4 Na_2_-ATP, 0.4 Na_3_-GTP, pH = 7.25, adjusted with KOH). The cells were monitored with a 40× Olympus water-immersion objective lens, a microscope (Olympus, BX51 WI) configured for dodt gradient contrast (DGC), and a camera (Andor, iXon3, Belfast, County antrim, UK). Data acquisition was conducted with a multiclamp 700B amplifier and a Digidata 1440A (Molecular Devices, San Jose, CA), which was controlled by Clampex 10. Data were analyzed using Clampfit 11.2.

### Neuronal survival and synapse formation assays

15 μg of protein from UTF- or CAR-astrocyte CM was added to neural media. 15000 neurons were plated and their survival was assessed after 3 days. To assay synapse formation, neurons were cultured for 7 days in normal neural media and for 6 more days in neural media with CM protein of UTF or CAR astrocytes. The cells were then fixed and stained for PSD95 and SYN. Puncta Analyzer plugin was used to quantify synapses in ImageJ.

### Secreted protein detection

15 μg of protein from UTF- or CAR-astrocytes (postnatal day 7) CM was applied to SDS-PAGE, and the antibodies against APOE, TSP1 and APP were used to detect these proteins by Western-blot, respectively.

### In vitro engulfment assay

Synaptosomes and crude CNS myelin were purified as described previously(33, 34). Synaptosomes/myelin were conjugated with pHrodo Red succinimidyl ester (Thermo Fisher Scientific, P36600) in 0.1 M sodium carbonate (pH 9.0) at room temperature for 2 h. Unbounded pHrodo was washed out by centrifugation and pHrodo-conjugated synaptosomes/myelin were re-suspended with isotonic buffer containing 5% DMSO for subsequent freezing. UTF or CAR astrocytes were incubated with 5 μl pHrodo-conjugated synaptosomes or 800 μg ml^-1^ medium pHrodo conjugated myelin debris for 24 h and analyzed by FACS.

### Western blot analysis

Protein samples from brain lysates and cell lysates were electrophoresed on 12% SDS-PAGE gels and transferred to polyvinylidene fluoride membranes. After blocking with 5% nonfat milk for 1 h at room temperature, the membrane was probed overnight at 4 °C with a primary antibody. After washing, HRP-conjugated secondary antibodies were used at a concentration of 1:5,000 at room temperature. The bands in immunoblots were visualized by enhanced chemiluminescence using an Amersham imager 680 imaging system (GE Healthcare) and quantified by densitometry and Image J software. Primary antibodies to the following proteins were used: RhoA (Invitrogen, #PA5-87403, 1:500), Rac1 (Invitrogen, #PA1-091X, 1:500), Cdc42 (Invitrogen, #PA1-092, 1:500), SOCS1 (Sangon, #D260748, 1:1000), SOCS3 (Sangon, #D121242, 1:1000), p65 (Cell Signaling Technology, #8242, 1:1000), MerTK (Abcam, #ab95925, 1:1000), pMerTK (Bioss, #bs18791R, 1:1000), IκB-α (Cell Signaling Technology, #4814, 1:1000), W20 (Monoclonal antibody, prepared by our laboratory, 1:500), Iba-1 (GeneTex, #GTX101495, 1:1000), GFAP (Cell Signaling Technology, #3670S, 1:1000); Clec7a (Invitrogen, #PA5-87072, 1:500), P2ry12 (Abcam, #ab188968, 1:500), PSD95 (Abcam, #ab12093, 1:1000), Synaptophysin (Abcam, #ab32127, 1:1000), APOE (Beyotime, #AF1921, 1:1000), TSP1(Beyotime, #AF8154, 1:1000) and APP (Abcam, #ab15272, 1:1000)

### Immunocytochemistry analysis

Cells cultured on glass slides were fixed with 4% PFA for 20 min at room temperature, blocked with block buffer (10% donkey serum in PBS + 0.3% Triton-X100) for 1 h, incubated with primary antibodies overnight at 4 °C, and then incubated with fluorescently conjugated secondary antibodies (Invitrogen) for 2 h at room temperature in dark and mounted on coverslips with antifade mounting medium (Solarbio, # S2110). Fluorescence signals were captured on a laser scanning confocal microscope (Leica TCS SP8). Primary antibodies against the following proteins were used: MerTK (Abcam, #ab95925, 1:100), W20 (Monoclonal antibody, prepared by our laboratory), Aβ (Biolegend, #803001, 1:100; Abcam, #ab201060), LAMP1 (Abcam, #ab24170, 1:100), Iba-1 (GeneTex, #GTX101495, 1:50); MAP2 (Abcam, #ab11267, 1:100), GFAP (Cell Signaling Technology, #3670S, 1:100; Abcam, #ab4674, 1:100).

### Immunohistochemistry analysis

Mice were deeply anaesthetized and immediately perfused with ice-old PBS containing heparin (10 U/mL) and sacrificed. Mouse brains were immediately removed and divided along the sagittal plane. The left brain hemisphere was fixed in 4% paraformaldehyde at 4 °C overnight, and 20-μm sagittal sections were then obtained using a Lecia CM1850 microtome. Before staining, the sections were incubated with sodium citrate buffer (10 mM sodium citrate, pH 6.0) for 20 min at 95 °C for antigen retrieval. The sections were next permeabilized and blocked with 10% normal donkey serum in 0.3% Triton X-100 PBST for 1 h at room temperature and then incubated with primary antibodies, followed by corresponding fluorescently conjugated secondary antibodies, respectively, and imaged on a Leica TCS SP8 confocal microscope. For 3’-Diaminobenzidine (DAB) immunostaining, the sections were incubated with a primary antibody, followed by a corresponding HRP-labeled secondary antibody and visualized with DAB by an Olympus IX73 inverted microscope with DP80 camera. All images were analyzed by Image J Software. Primary antibodies to the following proteins were used: W20 antibody (Monoclonal antibody, prepared by our laboratory), Iba-1 (Abcam, #ab178847, 1:100; GeneTex, #GTX101495, 1:50), GFAP (Cell Signaling Technology, #3670S, 1:100), Aβ (Biolegend, #803001, 1:100), PSD95 (Abcam, #ab12093 or #ab18258, 1:100); Synaptophysin (Abcam, #ab32127, 1:100).

### Animal and drug administration

In this study, all animal experiments were performed in accordance with the China Public Health Service Guide for the Care and Use of Laboratory Animals. Experiments involving mice and protocols were approved by the Institutional Animal Care and Use Committee of Tsinghua University (AP#15-LRT1). Six-month-old male APP/PS1 mice and their wild type littermates were obtained from Jackson Laboratory. APP/PS1 mice were treated with PBS (APP/PS1-PBS), empty liposomes (APP/PS1-NP), plasmid (1)-liposome lipoplexes (APP/PS1-CAR) and plasmid (2)-liposome lipoplexes (APP/PS1-ns-CAR). Wild type littermates treated with PBS (WT-PBS) and liposomes (WT-NP). The mice were intranasally administered with lipoplexes containing 1 μg plasmid or just PBS and empty liposomes every two days for 6 weeks. After the last administration, the behavioral tests were performed.

### Morris water maze (MWM) tests

Five days after the last administration, spatial memory of APP/PS1 mice was evaluated by Morris water maze tests, as described previously with minor modification^11^. Briefly, MWM tests were conducted in a circular 120 cm-diameter pool filled with opaque water kept at 21 °C and a platform (10 cm in diameter) submerged 1.0 cm under the water. On training days 1-5, the mice were allowed to swim for 60 s to find the platform, on which they were allowed to stay for 20 s. Mice unable to locate the platform were guided to it. The mice were trained twice a day over five consecutive days, with an inter-trial interval of 3 h to 4 h. At 24 h after the last learning trial, the mice were tested for memory retention in a probe trial without the platform. The swimming activity of each mouse was automatically recorded via a video tracking system using a video camera (Sony Corp.) mounted overhead.

### Y-maze test

The Y-maze test was conducted as previously described in detail(35). In brief, Y-maze testing consisted of 2 trials separated by an interval of 1 h. The first trial was 10 min in duration, the mouse was allowed to explore only 2 arms (the start and familiar arms) of the maze, and the third arm (novel arm) was blocked. In the second trial, the mice were put back in the same starting arm as in trial 1 with free access to all three arms for 5 min. By using a ceiling-mounted charge-coupled device camera, all trials were recorded on a videocassette recorder, and the number of entries and time spent in each arm in the video recordings were analyzed.

### Novel object recognition (NOR) test

The novel object recognition test was performed as described previously with some modifications(36, 37), Briefly, mice were individually habituated to explore the behavioral arena (50 cm × 50 cm × 25 cm white plastic box, empty) for 5 min one day before testing. Then, the mouse was removed from the box and placed back to its housing cage. During training test, each mouse was placed in the white box containing two identical objects for 5 min and returned quickly to its housing cage. The recognition memory was tested after 24 h. The time spent exploring and sniffing each object was recorded. The results are expressed as the discrimination index, which was calculated by subtracting the time spent exploring the familiar object from the time spent exploring the novel object and dividing by the total exploration time.

### Microdialysis

In vivo microdialysis sampling of brain ISF AβOs was performed in awake and behaving APP/PS1 mice as previously described(38). Briefly, guide cannulas (PEG-4; Eicom, Japan) were stereotaxically implanted above the left hippocampus, and microdialysis probes with a 3.0-mm, 1000-kDa-MWCO polyethylene membrane (PEP-4-03; Eicom) were inserted into the hippocampus through guide cannula. The probe and connecting tubes were perfused with an artificial cerebrospinal fluid perfusion buffer (in mM: 1.3 CaCl2, 1.2 MgSO4, 3 KCl, 0.4 KH2PO4, 25 NaHCO3, and 122 NaCl, pH 7.35) for 6 h at a flow rate of 1 μL min^-1^. Samples were collected at a flow rate of 0.5 μL min^-1^ and stored at 4°C in polypropylene tubes.

### Measurements for Aβ40, Aβ42 and AβOs

AβOs levels in the mouse brain homogenates of TBS-soluble fractions were measured using oligomeric Aβ ELISA kit (Biosensis, #BEK-2215-1P) according to the manufacturer’s instructions. The TBS-insoluble (guanidine-soluble) Aβ levels in the brain lysates of mice were quantified by ELISA using Aβ40 and Aβ42 immunoassay kits (IBL) following the manufacturer’s protocols. The levels of insoluble Aβ were standardized to the brain tissue weight and expressed in picogram or nanogram of Aβ per milligram of brain tissue.

### Measurements for inflammatory cytokines

For the inflammatory cytokine measurements, the levels of pro-inflammatory cytokines (IL-1β, TNF-α) and anti-inflammatory cytokines (TGF-β, IL-4) in samples from brain lysates of mice and cell culture supernatants were determined using corresponding ELISA kits (pro-inflammatory cytokine ELISA kits were obtained from Neobioscience technology, anti-inflammatory cytokine ELISA kits were obtained from Bioss) according to the manufacturer’s protocols. The absorbance at 450 nm was measured using a SpectraMax M5 microplate reader.

### Statistical analysis

Data were analyzed with GraphPad Prism v.8.0.1. Each figure legend denotes the statistical test used. All data are represented as mean ± s.e.m. unless otherwise indicated, Statistical significance was assessed using student’s *t*-test, one-way or two-way ANOVA followed by Tukey’s multiple comparisons test. In all cases, statistical difference was considered significant at *P < 0.05, **P < 0.01, ***P < 0.001. **** P < 0.0001

## Author contributions

R.-T.L. designed the experiments; L.Z., X.-G.L. and X.-Y.S. performed behavioral experiments; F.C., J.Z., X.-Y.D., X.-Y.N. and L.-J.L. performed primary microglial and astroglial cultures experiments. L.Z., X.-G.L., Y.-R.H. and C.-Y.L. performed immunocytochemistry and immunohistochemistry; L.Z. S.-J.H. and D.-Q.L. conducted the biochemistry experiments; K.W. and S.-Y.L. conducted western blot and qPCR experiments. H.A.R., L.S. and X.Q.W. performed electrophysiology experiments; J.-J.Y., X.-L.Y., S.-Y.J., W.-W.Z. and X.-X.X. provided samples and advice; L.Z., S.L., X.-G.L. and D.-Q.L. analyzed the data; R.-T.L. and L.Z. wrote the manuscript.

---

**Fig. S1.**
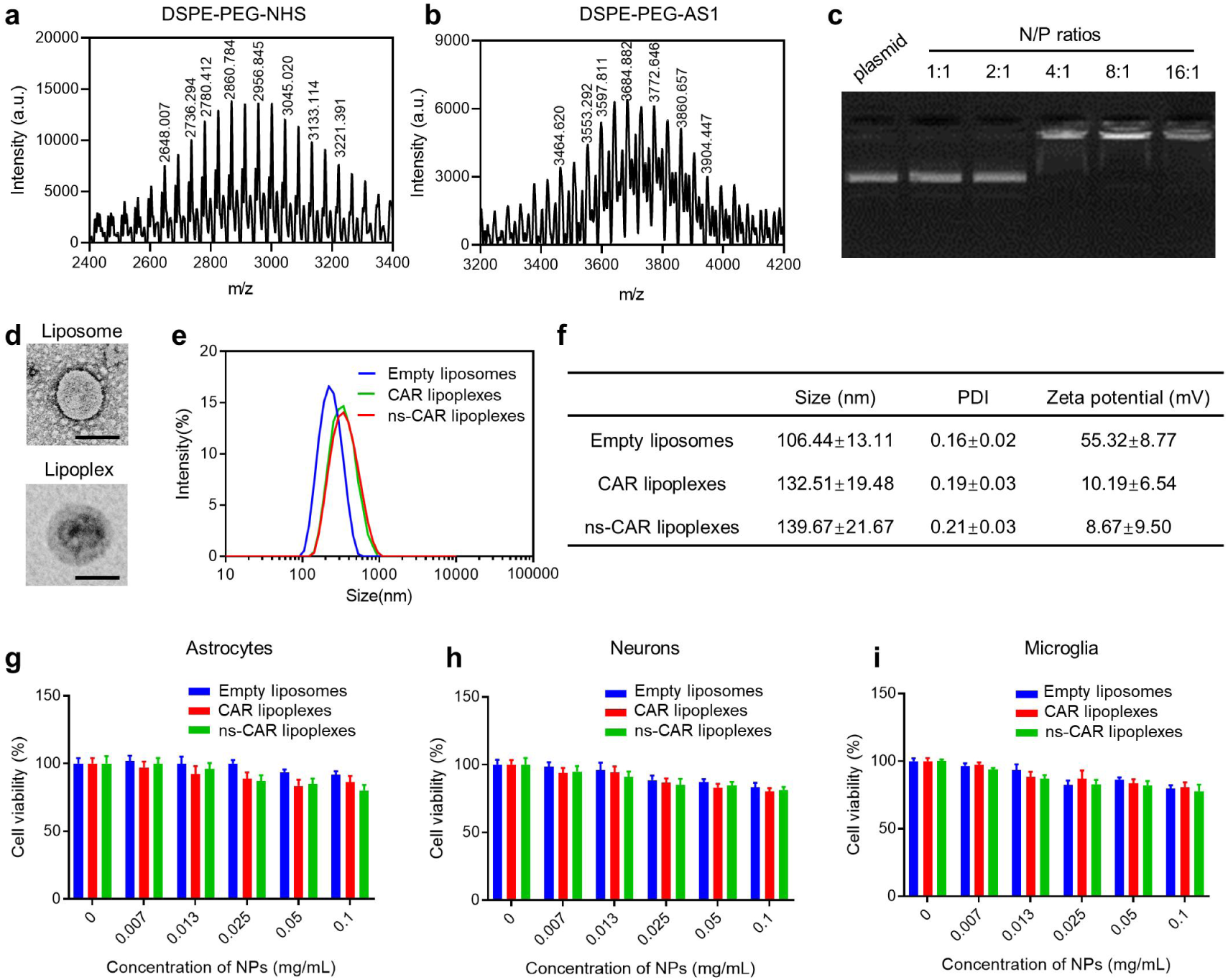
Characterization of the prepared lipoplexes. **a-b**, The MALDI-TOF-mass spectrum of DSPE-PEG-NHS **(a)** and DSPE-PEG-AS1 **(b)**. **c**, The N/P ratio of plasmid-liposome lipoplexes measured by gel retardation assay. **d**, Transmission electron micrographs of a representative liposome (up) and lipoplex (down). Scale bars: 100 nm. **e**, Representative size distributions of empty liposomes, CAR lipoplexes and ns-CAR lipoplexes measured by dynamic light scattering method. **f**, Table of size, PDI and zeta potential for empty liposomes, CAR lipoplexes and ns-CAR lipoplexes. Data are represented as mean ± s.e.m. of n = 3 technical replicates. **g-i**, Toxicity of empty liposomes, CAR lipoplexes and ns-CAR lipoplexes on astrocytes, neurons and microglia. Different concentrations of empty liposomes, CAR lipoplexes and ns-CAR lipoplexes were added to the culture of primary astrocytes, neurons and microglia, respectively. Cell viability was measured by MTT analysis after 72 h-incubation. Data is represented as mean ± s.e.m. of n = 6 technical replicates. Experiment was repeated at least 3 times.

**Fig. S2.**
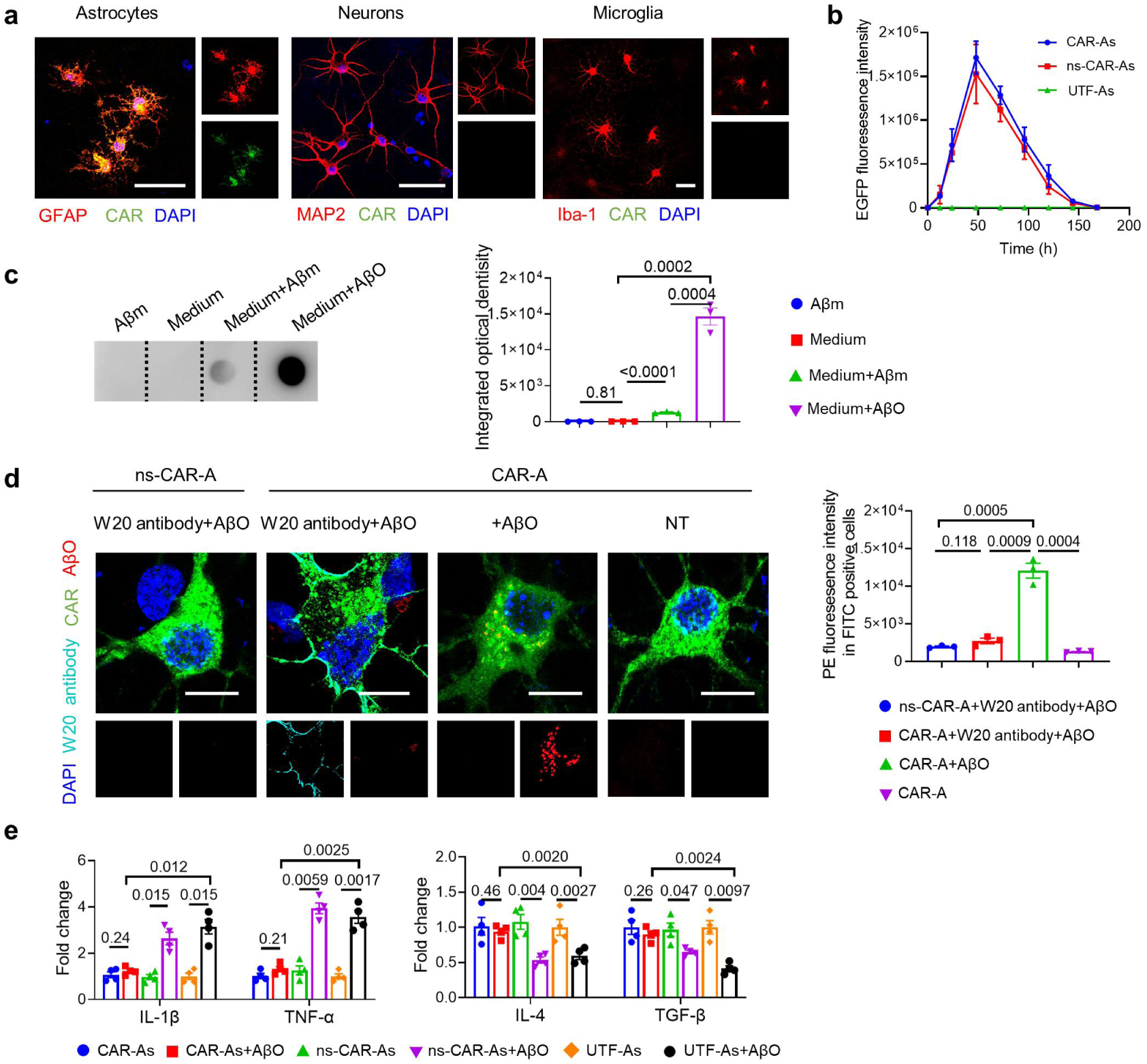
Expression of CARs and their effect on AβOs engulfment and inflammatory generation. **a**, Representative images depicting the expression of CARs in astrocytes (left), neurons (middle), microglia (right). Scale bars: 50 μm. **b**, Flow cytometric quantification of the EGFP fluorescence signal of astrocytes 0-168 h after transfection with CAR lipoplexes and ns-CAR lipoplexes. **c**, Left: 200 nM Aβ monomers or Aβ oligomers were added to medium and were incubated at 37 °C for 6 h, the amount of oligomers in medium were immunoblotted by W20. Right: the integrated optical density was quantified using Image J pro. **d**, Left: anti-W20 antibody blocked AβOs engulfment by CAR-As. CAR-As were incubated with or without anti-W20 antibody for 2 h, then AβOs were added and incubated for another 6 h. Scale bars: 5 μm. Right: the amount of AβOs in CAR-As were analyzed by flow cytometry. **e**, The levels of secreted pro-inflammatory cytokines (IL-1β, TNF-α) and anti-inflammatory cytokines (IL-4, TGF-β) determined by ELISA. Data are represented as mean ± s.e.m. of n = 3 technical replicates and are representative of four experiments. Statistical significances were determined by one-way ANOVA with Tukey’s multiple comparisons test.

**Fig. S3.**
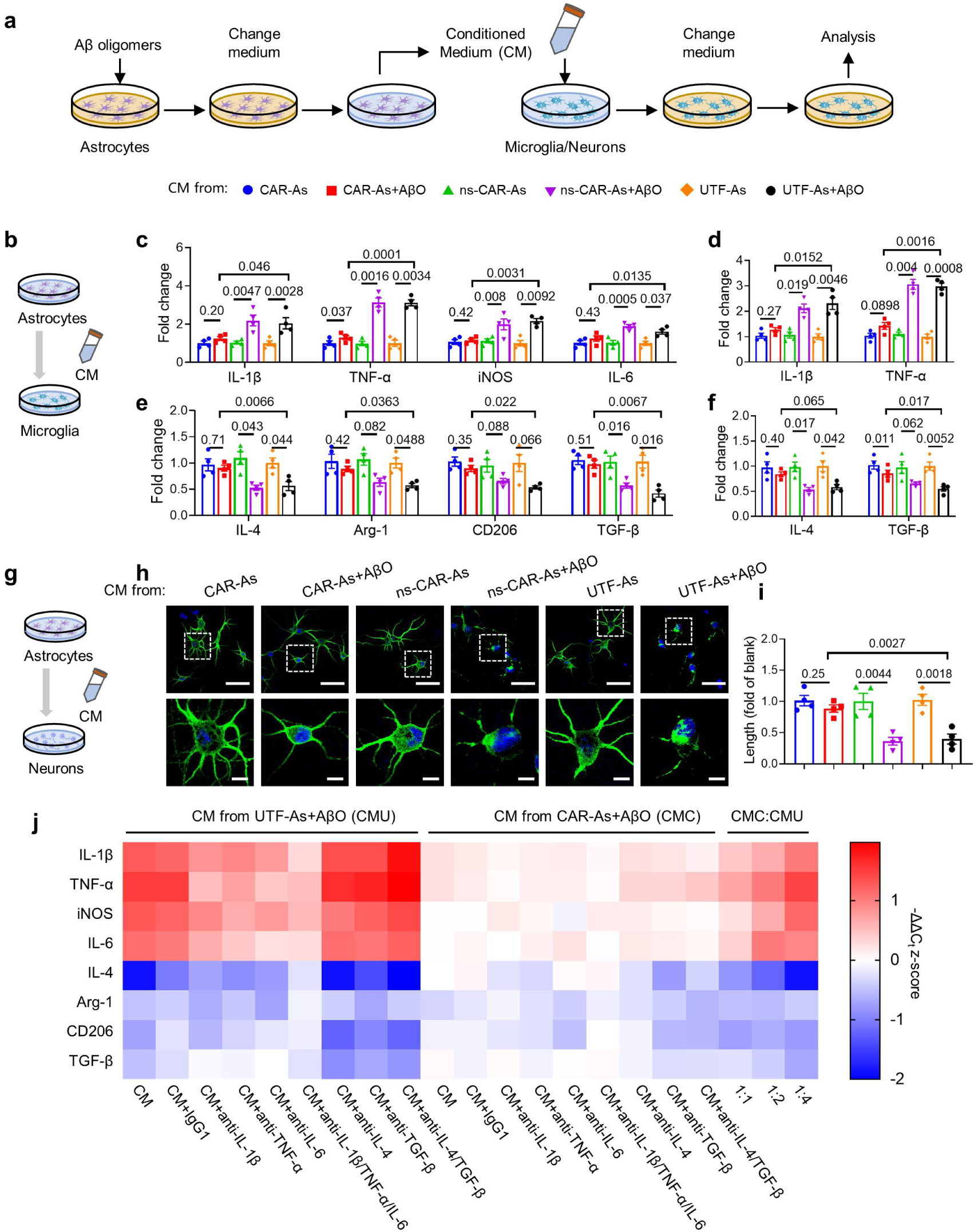
The effect of CAR-As on microenvironment, microglia and neurons. **a**, Cartoon depicting the experimental paradigm for identifying the microenvironmental influences in microglia and neurons by CAR-As using conditional medium (CM). **b-f**, CAR-As, ns-CAR-As or UTF-As treated with or without AβOs for 24 h, after replacing the medium and culturing for another 24 h, the CM were added to microglia culture (**b**), after 24 h-incubation, the pro-inflammatory cytokines (IL-1β, TNF-α, iNOS, IL-6) (**c**) and anti-inflammatory cytokines (IL-4, Arg-1, CD206, TGF-β) (**e**) in the microglia were determined by RT-qPCR. The secreted pro-inflammatory cytokines (IL-1β, TNF-α) (**d**) and anti-inflammatory cytokines (IL-4, TGF-β) (**f**) in the medium were determined by ELISA. Data are represented as mean ± s.e.m. of n = 3 technical replicates and are representative of four experiments. Statistical significance was calculated with ANOVA with multiple comparisons. **g-i**, CAR-As, ns-CAR-As or UTF-As treated with or without AβOs for 24 h, after replacing the medium and culturing for another 24 h, the CM were added to neurons culture (**g**), after 24 h-incubation, the morphology of the neurons was imaged by confocal microscopy (**h**). The profiles shown here are representative of three independent experiments. Scale bars: 50 μm (up), 10 μm (down). The length of axons and dendrites were quantified using Image J pro (**i**). Values were normalized against the blank group. Data are represented as mean ± s.e.m. of n = 3 technical replicates and are representative of four experiments. Statistical significances were determined by one-way ANOVA with Tukey’s multiple comparisons test. **j**, Heat map of pro-inflammatory and anti-inflammatory cytokine transcripts in microglia. Anti-IL-1β, anti-TNF-α, anti-IL-6, anti-IL-10 and anti-TGF-β neutralizing antibodies were added to the CM from UTF-A+AβO or CAR-As+AβO groups, respectively and the CM with different neutralizing antibodies were added to microglia culture, after 24 h-incubation, the transcription of pro-inflammatory cytokines and anti-inflammatory cytokines were determined by RT-qPCR.

**Fig. S4.**
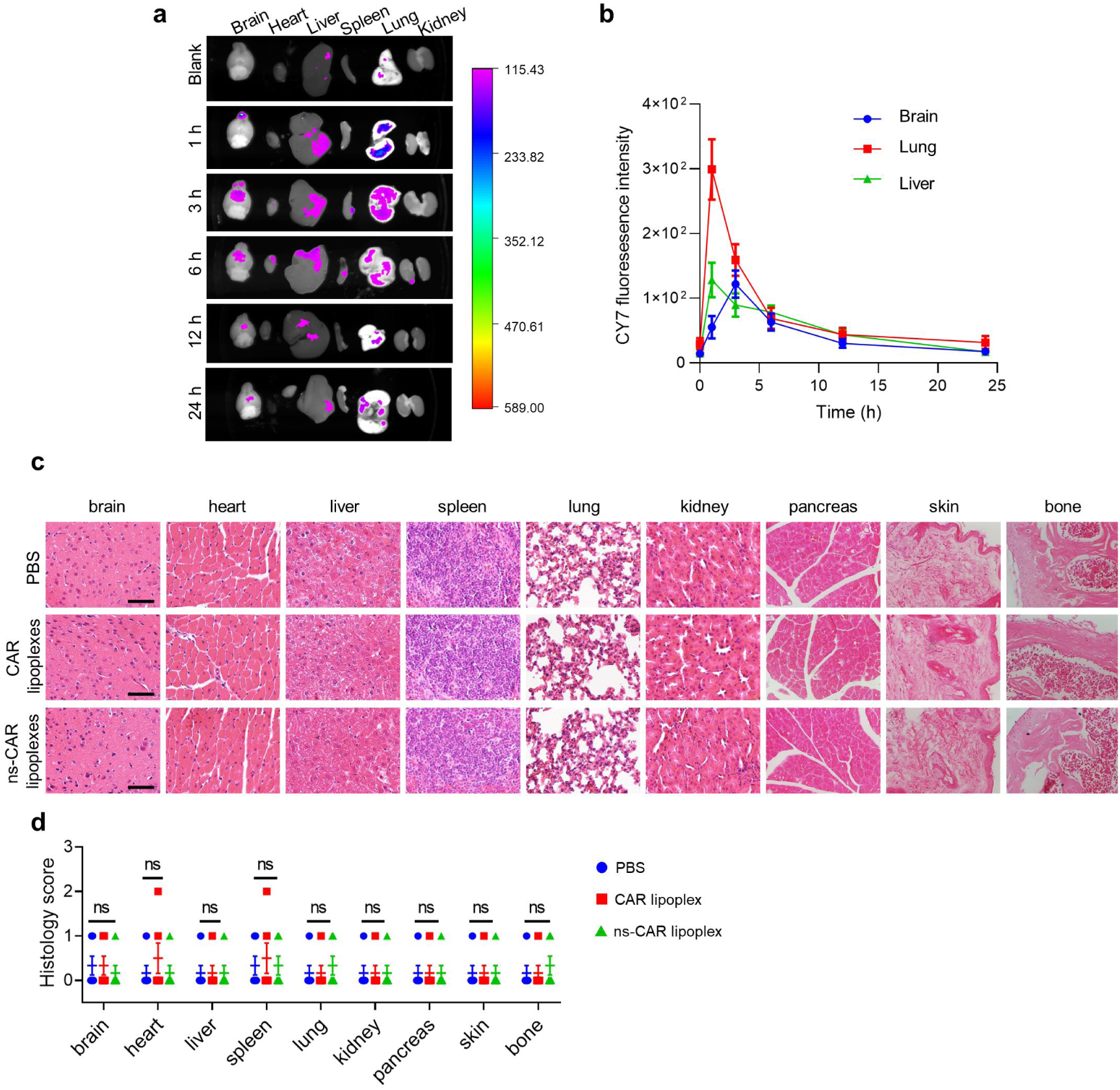
Distribution and biosafety of CAR lipoplexes in vivo. **a**, The distribution of CAR lipoplexes in mice. The mice were intranasally treated with lipoplexes labeled with CY-7. The fluorescence in various organs was detected using IVIS spectrum imaging system at different time points. **b**, Mean concentration-time profiles of CY-7 in brain, lung and liver after intranasal administration of CAR lipoplexes. **c**, The biosafety of lipoplexes. The treated mice were sacrificed and the organs and tissues were harvested at 6 weeks after the injection. The sections of brain, heart, liver, spleen, lung, kidneys, pancreas, skin and bone were stained with H&E . Scale bars: 50 μm. d, The histology scores of different tissues were determined according to INHAND (0 indicates no change and 1–4 indicate increasing severity).

**Fig. S5.**
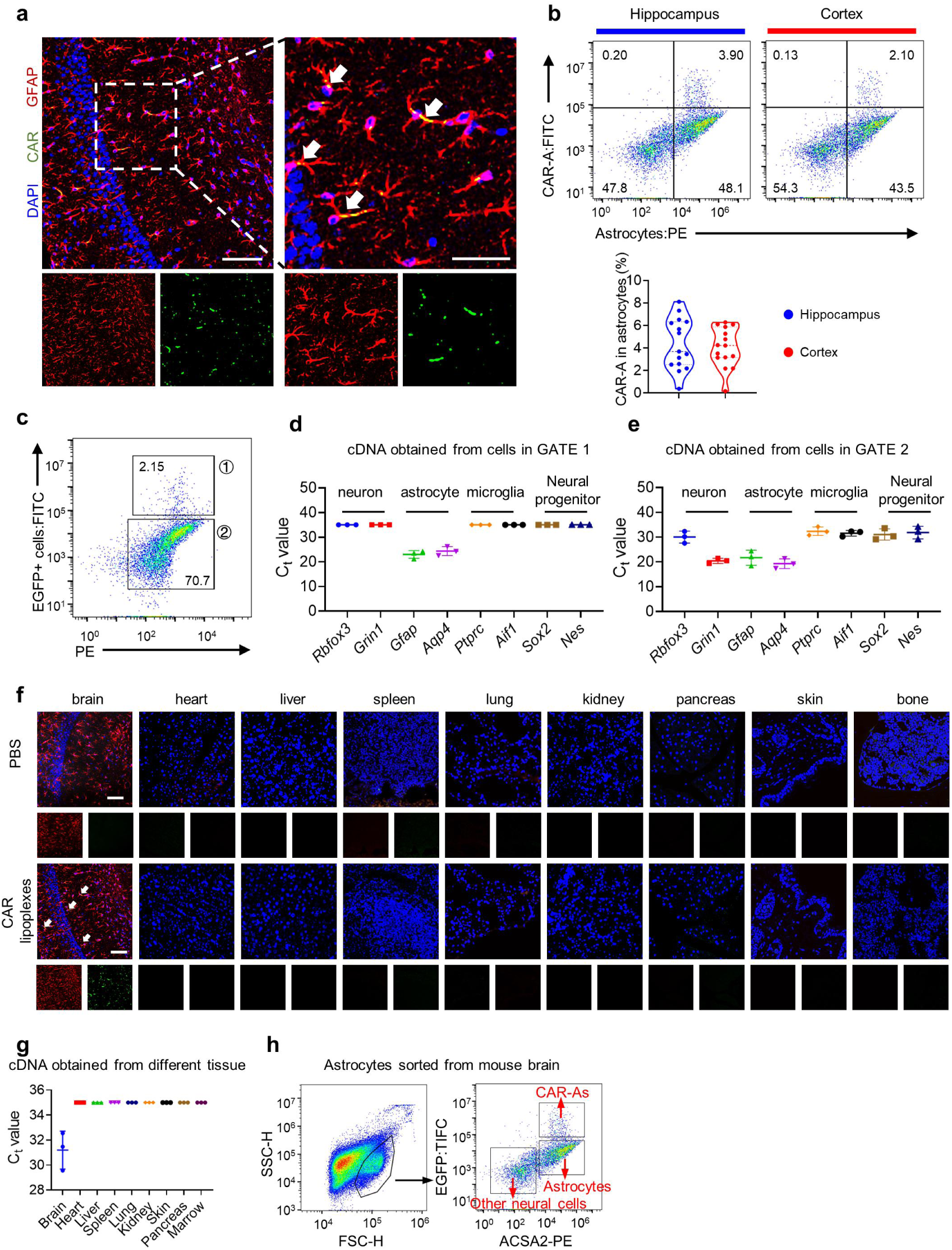
CARs specifically expressed in astrocytes in mouse brains. **a**, Left: representative images showing the CARs expressed in astrocytes in the mouse brains treated with CAR lipoplexes 72 h later. Scale bar: 100 μm. Right: enlarged colocalization images from the white dashed box in left image. Scale bar: 50 μm. **b**, Up: flow cytometry analysis of transfection efficiency of CAR lipoplexes in hippocampus and cortex. Down: the EGFP positive astrocytes (CAR-As) in total astrocytes were quantified. **c**, Sorting scheme for isolation of CAR^+^ cells and CAR^-^ cells in mouse brains. **d-e**, Identification of cell type of CAR^+^ cells and CAR^-^ cells by qPCR. cDNA was obtained from CAR^+^ cells and CAR^-^ cells, followed by qPCR tests for specific markers for neurons, astrocytes, microglia and neural progenitors. **f**, Representative images showing the expression of CARs in various organs of mice treated with CAR lipoplexes 72 h later. Red for astrocytes, green for CARs or ns-CARs and blue for nuclei. Scale bars: 100 μm. **g**, CAR expression in various organs were detected by qPCR. cDNA was obtained from various organs, followed by qPCR tests for specific markers for CAR. **h**, Sorting scheme for isolation of CAR-As and astrocytes from mouse brain.

**Fig. S6.**
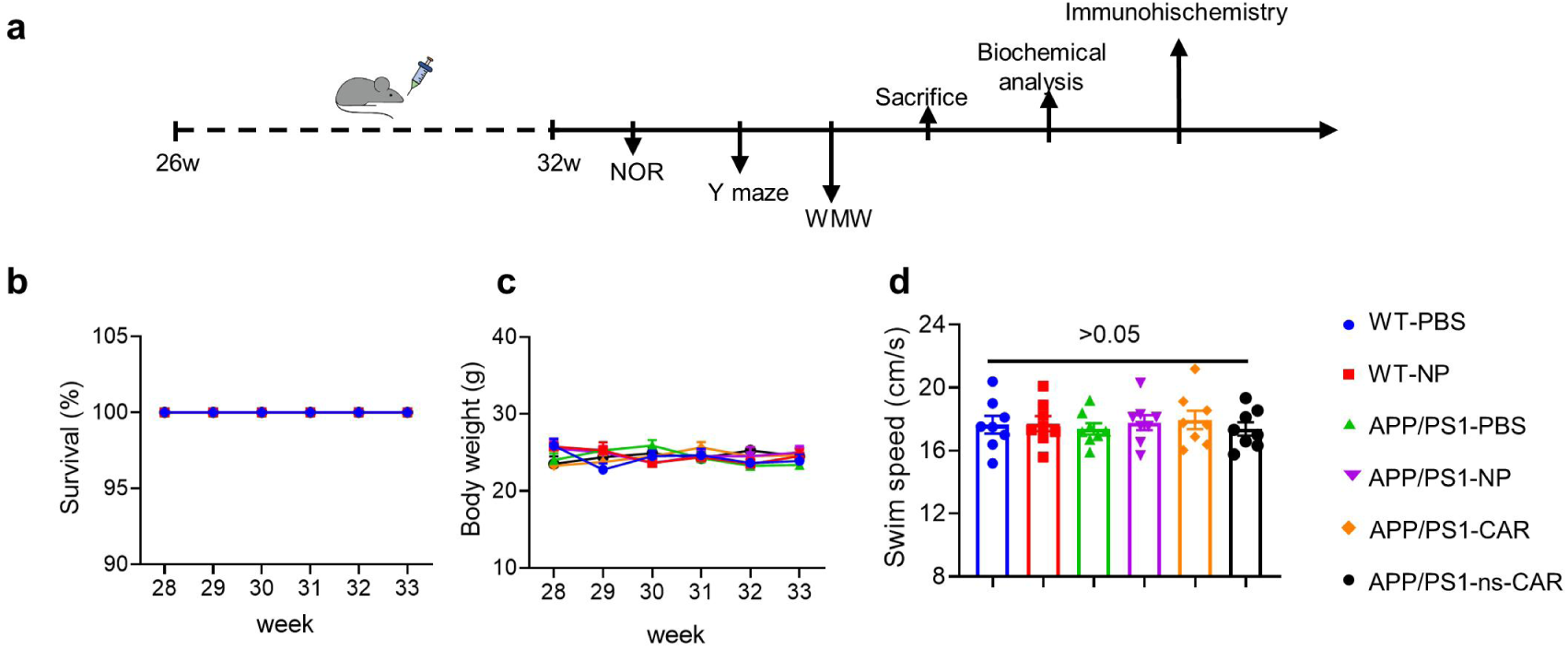
The effect of CAR-As on survival, body weight, swim speed of APP/PS1 mice. **a**, Schematic diagram of intranasal treatment, behavioral test and the physiological and biochemical analysis for APP/PS1 mice. **b-d**, Wild type mice were intranasally treated with saline and empty liposomes, and APP/PS1 mice were intranasally treated with saline, empty liposomes, CAR lipoplexes and ns-CAR lipoplexes for 6 weeks, the mouse survival, body weight and swim speed were consecutively detected within 28-33 weeks. For panel **b-d**, Values are the mean ± s.e.m. of n = 6-8 mice per group. A two-way ANOVA with post hoc Tukey test was used for statistical analysis.

**Fig. S7.**
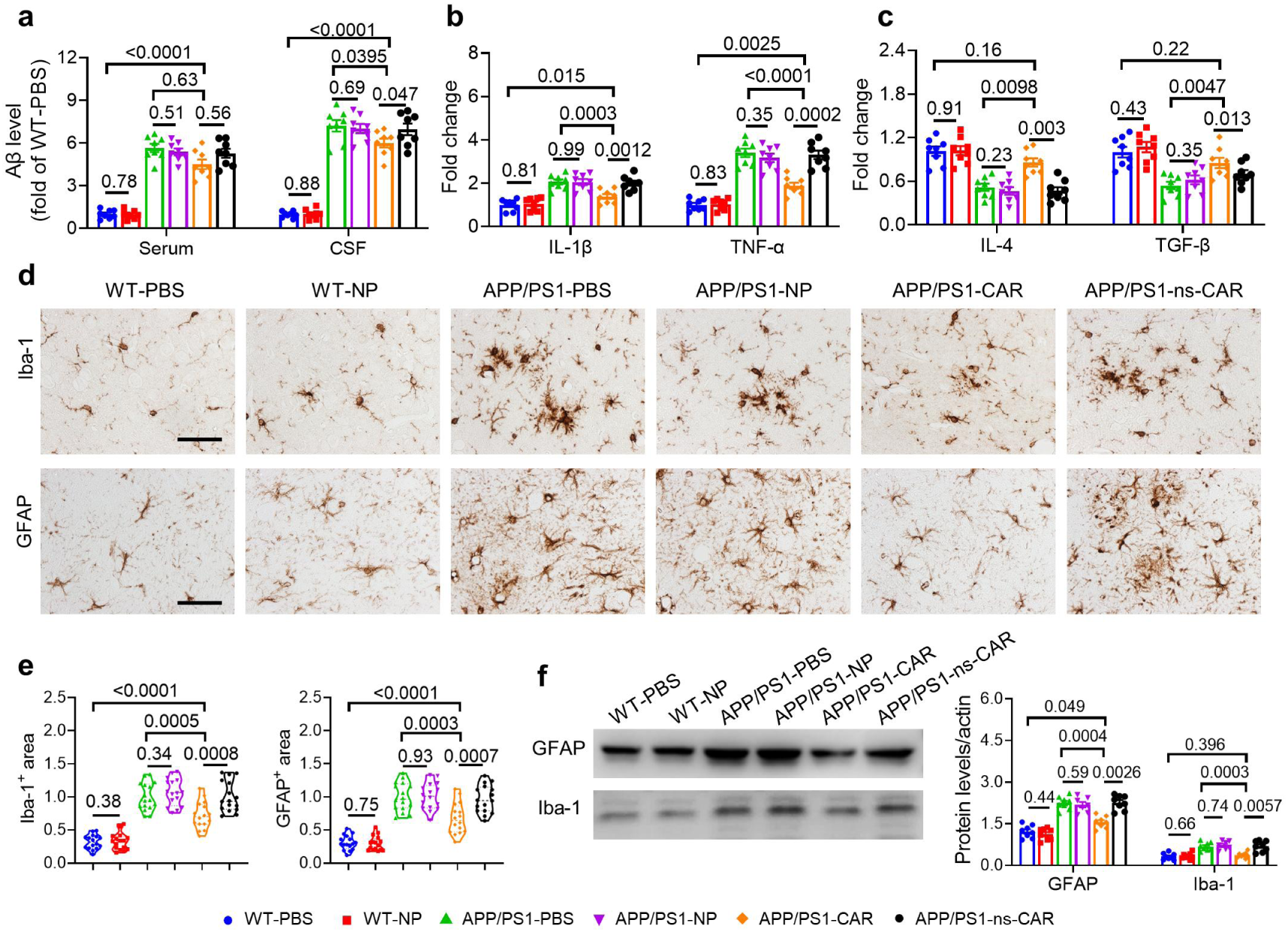
CAR-As ameliorated the neuropathology in the brains of APP/PS1 mice. Wild type mice were treated with saline and empty liposomes, and APP/PS1 mice were intranasally treated with saline, empty liposomes, CAR lipoplexes and ns-CAR lipoplexes for 6 weeks. **a**, Aβ levels in mouse serum and cerebrospinal fluid (CSF) in different experimental groups were measured by ELISA. **b-c**, The protein levels of pro-inflammatory cytokines (IL-1β, TNF-α) (**b**) and anti-inflammatory cytokines (IL-4, TGF-β) (**c**) in mouse brains were determined by ELISA. **d**, Representative images of microgliosis (up) and astrogliosis (down) in mouse brains in different experimental groups. Scale bars: 50 μm. **e**, The area of microglia and astrocytes in cortex and hippocampus were quantified using Image J pro. Values were normalized against the APP/PS1-PBS group. **f**, The protein levels of GFAP and Iba-1 were measured by Western blot (left), and quantified using Image J pro (right). Statistical significances were determined by one-way ANOVA with Tukey’s multiple comparisons test.

**Fig. S8.**
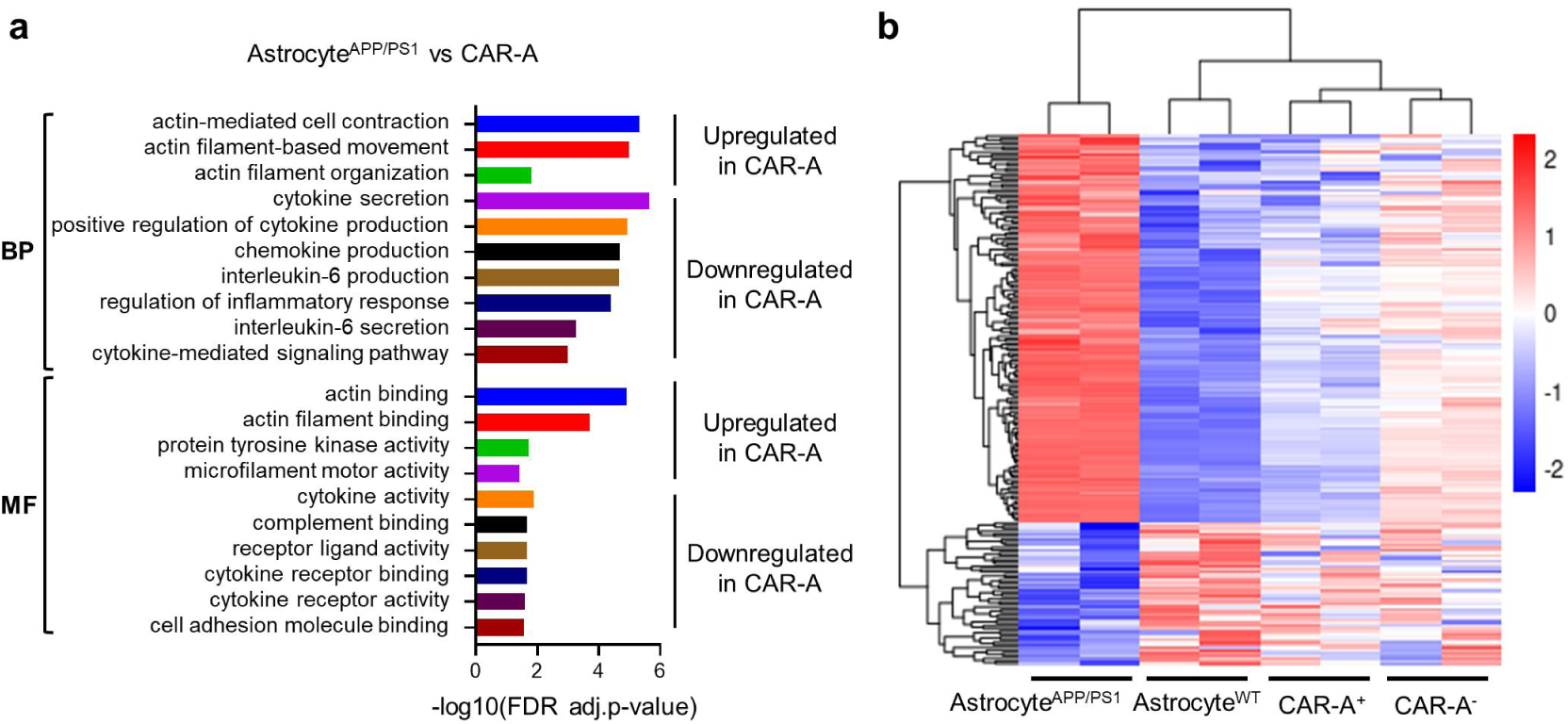
Comparison of gene expression profiles from astrocyte^WT^, astrocyte^APP/PS1^, CAR-A^+^ and CAR-A^-^. **a**, Transcriptomic GO analyses between the astrocytes^APP/PS1^ and CAR-As were performed with the FDR-adjusted p-value<0.05 (adjustments were made for multiple comparisons; FDR-corrected by Toppgene analysis). BP, biological process, MF, molecular function. **b**, RNA-seq analysis of differential gene expression between astrocytes^APP/PS1^, astrocytes^WT^, CAR-A^+^ and CAR-A^-^. n = 2 per group.

**Fig. S9.**
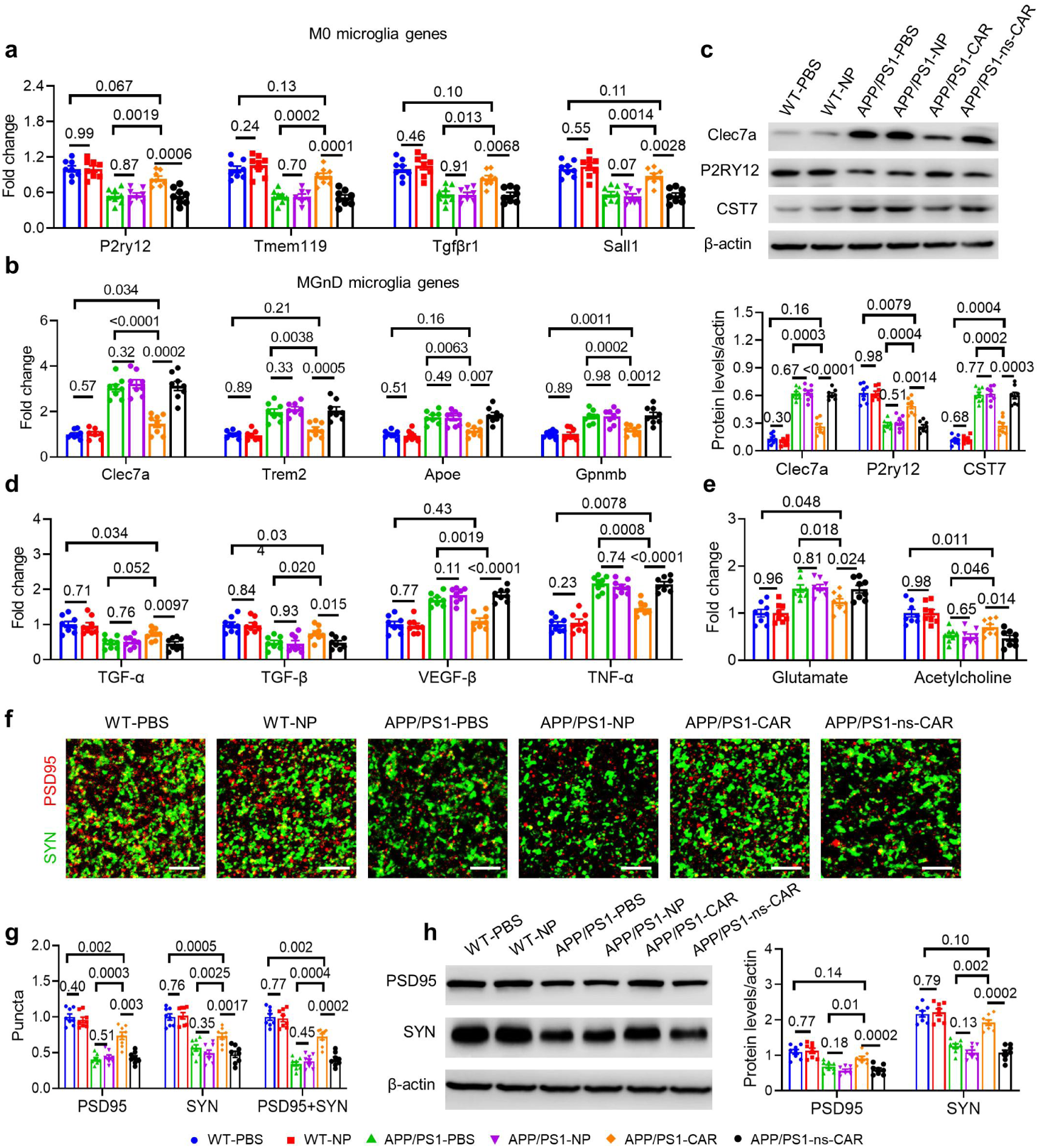
The effect of CAR-As on microglial phenotype and synapse level. Wild type mice were treated with saline and empty liposomes, and APP/PS1 mice were intranasally treated with saline, empty liposomes, CAR lipoplexes and ns-CAR lipoplexes for 6 weeks. **a-b**, Microglia were isolated from the mouse brains in different experimental groups, M0 microglia genes (P2ry12, Tmem119, Tgfβr1, and Sall1) (**a**) and MGnD microglia genes (Clec7a, Trem2, Apoe, and Gpnmb) (**b**) were determined by RT-qPCR, values were normalized against the WT-PBS group. Data are represented as the mean ± s.e.m. of n = 6-8 mice per group. A one-way ANOVA with post hoc Tukey test was used for statistical analysis. **c**, The protein levels of Clec7a, P2ry12 and CST7 were measured by Western blot, and quantified using Image J pro. **d**, TGF-α, TGF-β, VEGF-β and TNF-α were determined by RT-qPCR, values were normalized against the WT-PBS group. **e**, levels of glutamate and acetylcholine in mouse brain in different experimental groups were measured by ELISA. **f**, Representative images depicting level of synapses by staining the brain slice with anti-PSD95 and anti-SYN antibody. The puncta of co-localization of PSD95 and SYN indicated the intact synapse. Scale bars: 5 μm. **g**, The levels of PSD95, SYN and intact synapse were quantified using Image J pro. **h**, The protein levels of PSD95 and SYN were measured by Western blot (left), and quantified using Image J pro (right). Values were normalized against the WT-PBS group. Data are represented as the mean ± s.e.m. of n = 6-8 mice per group. A one-way ANOVA with post hoc Tukey test was used for statistical analysis.

**Fig. S10.**
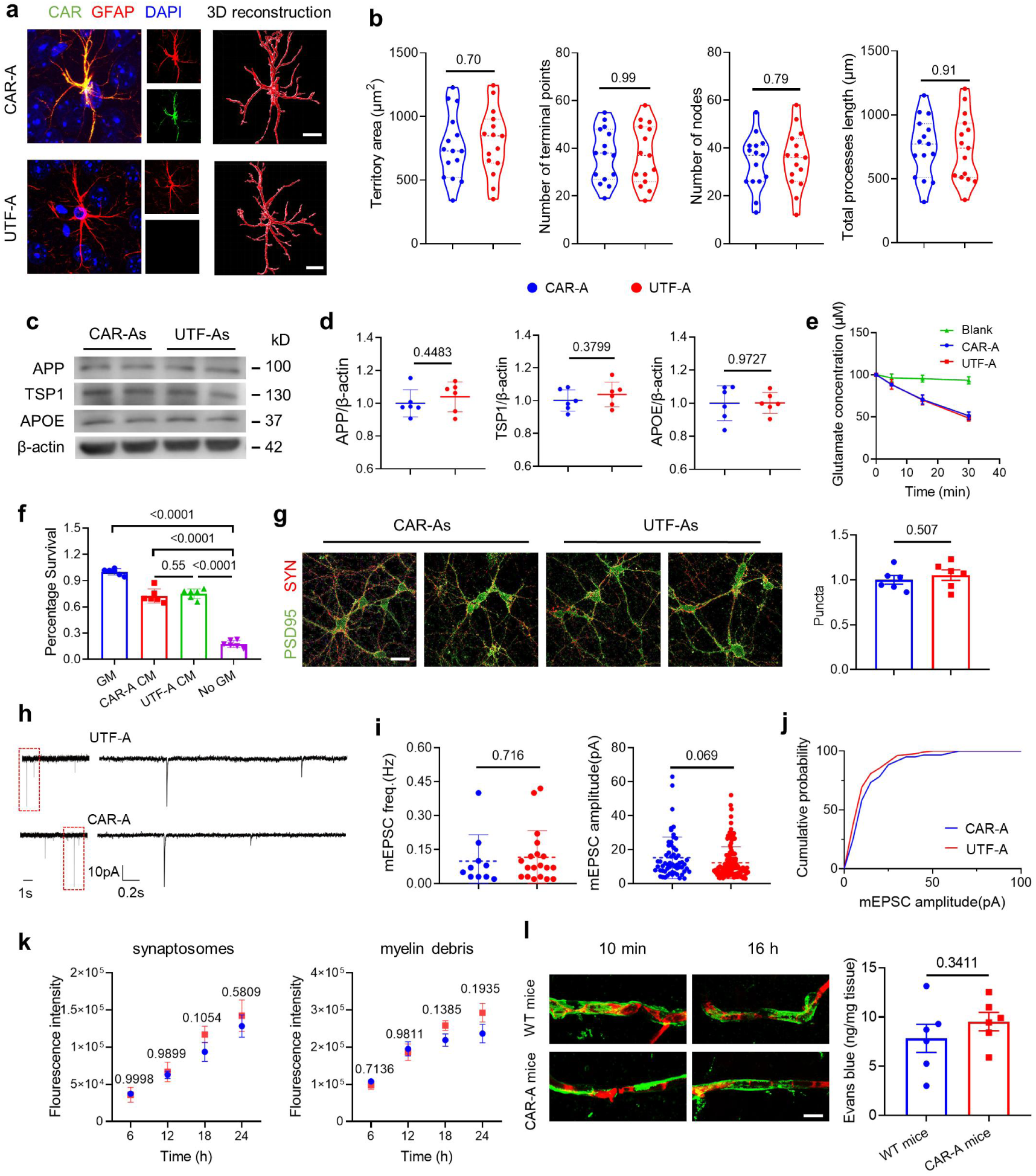
The morphology and major biological functions of CAR-As. **a**, Confocal micrographs of CAR-A and control astrocytes and 3D reconstructions of their soma and processes. Scale bars: 10 μm. **b**, Morphological quantification of CAR-As and astrocytes in the projected territory area, number of nodes, number of terminal points and total length of processes. **c,** Levels of APP, TSP1 and APOE produced by CAR-As and UTF-As. APP, TSP1 and APOE from cultured CAR-As and UTF-As were analyzed by Western blot, β-actin was used as controls. **d**, Relative levels of APP, TSP1 and APOE quantified using Image J pro. The results were combined to express as mean ± s.e.m of six independent repeats. **e**, Analysis of glutamate uptake in CAR-As, ns-CAR-As and UTF-As. Glutamate concentrations in media were measured after the media containing 100 μM glutamate were incubated with CAR-As, ns-CAR-As and UTF-As for 0∼30 min. **f,** The effect of conditioned media (CM) from CAR-As and UTF-As on neuronal survival. CM were added to neuron culture and the neuronal survival was detected, RGC growth media was used as controls. **g**, The effect of CM from CAR-As and UTF-As on neuronal synaptogenesis. Left: Confocal micrographs of neuron cultured with CM from CAR-As or UTF-As. Right: Synaptogenesis was quantified by assessing colocalization of presynaptic marker SYN (red) and postsynaptic marker PSD95 (green) with Image J. **h**, The representative traces of mEPSCs from neuron cultured with CM from CAR-As and UTF-As. **i**, Frequency (UTF-A: 0.11 ±0.03 Hz, CAR-A: 0.1 ±0.03 Hz) and amplitude (UTF-A: 12.26 ±0.82pA, CAR-A: 15.21 ± 1.56pA) of mEPSC, data shown as mean ± SEM. **j**, Cumulative amplitude of mEPSCs of neuron cultured with CAR-A CM or UTF-A CM. **k**, UTF-A or CAR-A were incubated with 5 μl pHrodo-conjugated synaptosomes or 800 μg ml^−1^ medium pHrodo-conjugated myelin debris for 24 h and analyzed by FACS. **l**, The effect of CAR-A on the infiltration of blood brain barrier. Left, the distribution of Evans blue (EB). Perivascular marker isolectin B4 (Ib4) in the mouse brains treated with CAR or UTF was stained via IHC 10 min or 16 h post EB injection and prior to perfusion. Scale bars: 25 μm. Right, EB fluorescence of brain homogenates obtained by a microplate reader after EB dye circulating for 16 h followed by 5-min perfusion.

**Fig. S11.**
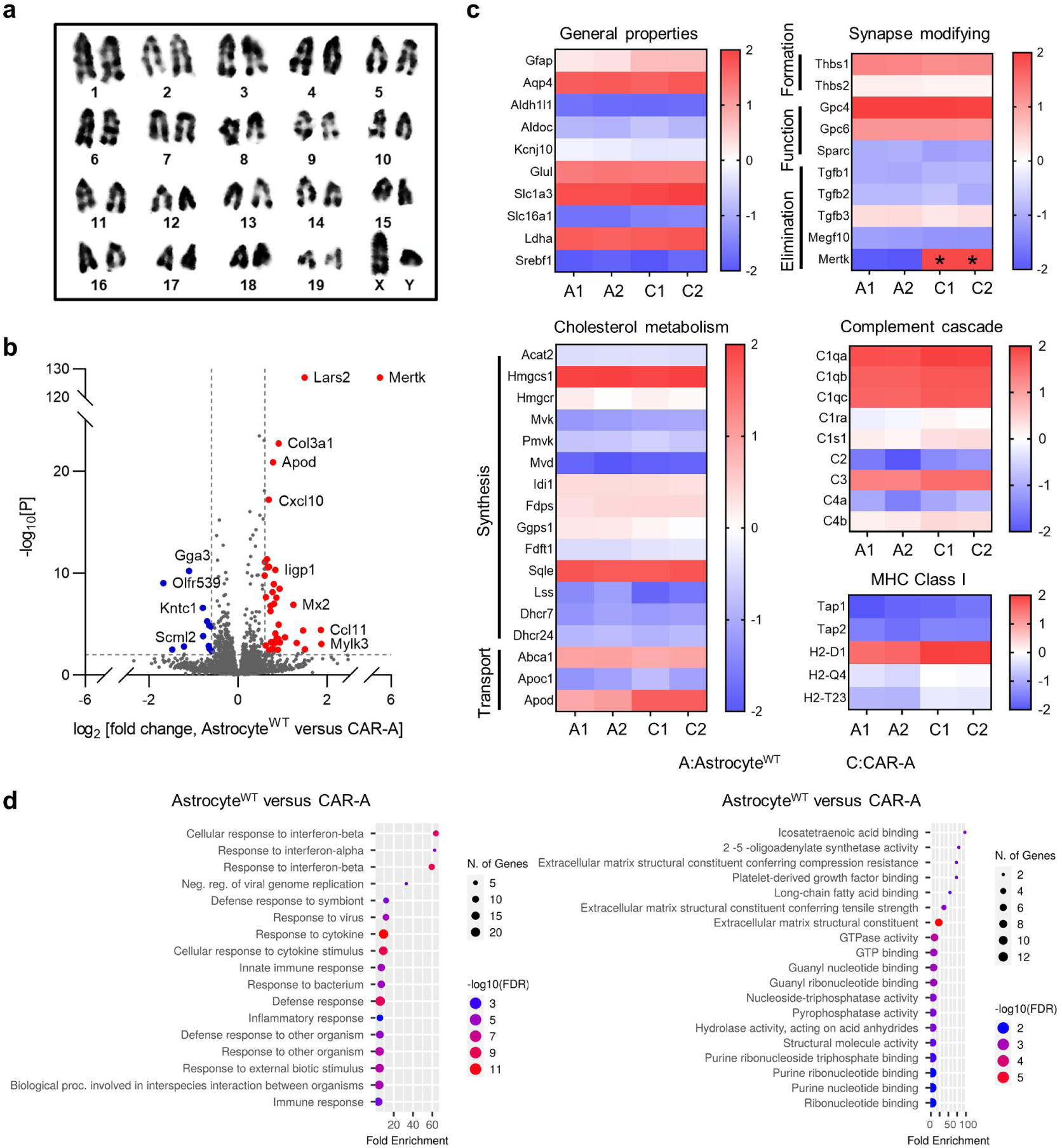
Differential transcriptome profiles between astrocyte^WT^ and CAR-A^+^. **a**, Karyotype of CAR-As. **b**, Volcano plot of differentially regulated genes from astrocyte^WT^ and CAR-A (FC>1.5, P<0.005). **c**, Heatmaps demonstrating the expression of genes important for astrocyte identity and function. The genes are mainly involved in cholesterol metabolism, immune/antigenic responses, and regulated neuronal synapse formation, function, and elimination in astrocyte^WT^ and CAR-A. **d**, Gene set enrichment analysis for astrocyte^WT^ and CAR-A group.

